# All Quiet on the Western Front? The evolutionary history of monogeneans (Dactylogyridae: *Cichlidogyrus, Onchobdella*) infecting a West and Central African tribe of cichlid fishes (Chromidotilapiini)

**DOI:** 10.1101/2023.01.23.525183

**Authors:** Tanisha Moons, Nikol Kmentová, Antoine Pariselle, Tom Artois, Wim Bert, Maarten P. M. Vanhove, Armando J. Cruz-Laufer

## Abstract

Owing to the largely unexplored diversity of metazoan parasites, their speciation mechanisms and the circumstances under which such speciation occurs—in allopatry or sympatry—remain vastly understudied. Cichlids and their monogenean flatworm parasites have previously served as study system for macroevolutionary processes, e.g. for the role of East African host radiations on parasite communities. Here, we investigate the diversity and evolution of the poorly explored monogeneans infecting a West and Central African lineage of cichlid fishes: Chromidotilapiini Greenwood, 1987, which is the most species-rich tribe of cichlids in this region. We screened gills of 149 host specimens (27 species) from natural history collections and measured systematically informative characters of the sclerotised attachment and reproductive organs of the parasites. Ten monogenean species (Dactylogyridae: *Cichlidogyrus* and *Onchobdella*) were found, eight of which are described and one redescribed herein. The phylogenetic positions of chromidotilapiines-infecting species of *Cichlidogyrus* were inferred through a parsimony analysis of the morphological characters. Furthermore, we employed machine learning algorithms to detect morphological features associated with the main lineages of *Cichlidogyrus*. Although the results of these experimental algorithms remain inconclusive, the parsimony analysis indicates that West and Central African lineages of *Cichlidogyrus* and *Onchobdella* are monophyletic, unlike the paraphyletic host lineages. Several instances of host sharing suggest occurrences of intra-host speciation (sympatry) and host switching (allopatry). Some morphological variation was recorded that may also indicate the presence of species complexes. We conclude that collection material can provide important insights on parasite evolution despite the lack of well-preserved DNA material.

**Data availability statement:** Type material was deposited in the invertebrate collection of the Royal Museum for Central Africa (Tervuren, Belgium) (RMCA) (RMCA_VERMES_XXXXX–XXX), the collection of the Research Group Zoology: Biodiversity and Toxicology of Hasselt University (Diepenbeek, Belgium) (HU XXX–XXX), the Finnish Museum of Natural History (Helsinki, Finland) (MZH XXXXX–XXX), and the Iziko South African Museum (Cape Town, South Africa) (SAMC-XXXXXXX-XXX). The morphological data that support the findings of this study are openly available in MorphoBank at www.morphobank.org, at https://dx.doi.org/XXXXXXXX. Phylogenetic trees and data matrices for the analysis in TNT are included as additional data in MorphoBank.

## Introduction

### Parasite speciation: sympatry or allopatry?

Parasitism is one of the most successful lifestyles in terms of extant species richness [89] and biomass [51]. Evolutionary processes in these organisms have recently received increased attention, especially in the context of emerging infectious diseases (see [8]). Yet, most evolutionary processes in parasites remain poorly investigated. This limited knowledge stems from our poor understanding of the evolutionary history of most parasite taxa, with the majority of species remaining undescribed [39,89,91] or lacking DNA sequence data [91]. Despite these knowledge gaps, the lifestyle of parasites predicts several distinct characteristics different from most free-living animals such as highly specialized feeding behaviour and shorter generation times [38]. Parasite populations are also often larger in numbers but also more fragmented than populations of free-living organisms [20, 38]. These characteristics influence evolutionary rates and effective population size [38]. Parasites have a strong potential for speciation [13,90,95,116], which arises from their narrow habitat selection that can be limited to single host species or even distinct microhabitats on a host species [26, 64], and the evolutionary arms races between hosts and parasites [45]. In some cases, these factors can lead to adaptive radiations (i.e. explosive species formation), of which some of the most spectacular examples are displayed by parasites including anisakid nematodes [50] and endoparasitic snails that infect corals [25].

As with many other organisms, parasite speciation can occur in allopatry or sympatry [38,64,116]. The major criterion in distinguishing between allopatric and sympatric speciation is whether the barrier to gene flow is extrinsic or intrinsic [24]. Extrinsic barriers prevent the mating between parasites from different host species because of geography or vector specificity [7, 24]. When applying the concepts of the ‘island hypothesis’ to parasites, host species and even individual hosts may be considered as the equivalent of separated habitat patches (‘islands’) for free-living organisms. The physical and/or phylogenetic distance between host species and/or individuals determines the permeability of the barriers separating these habitats [64]. Parasite speciation can, therefore, be a direct consequence of host speciation (*co-speciation*) or occur as *accidental host switching* from one species to another (also referred to as *lateral transfer*), which represents a form of geographic isolation of small populations [38]. Sympatric speciation in parasites occurs in the absence of physical barriers but in the presence of intrinsic barriers, e.g. different preferences for mating habitats and allelic incompatibilities within parasites infecting the same host species or individuals [70] (also referred to as *intra-host speciation*, *duplication*, or *synxenic speciation*). These barriers can result in the evolution of a sexual preference for individuals infecting the same host species [24, 64]. Sympatric speciation has also been suggested to result from particular ecological conditions that facilitate *host switching through active host selection* [35, 47] (not to be confused with *accidental host switching*, which is a passive mechanism, see above) such as the nutritional value of the host species, intensity of competition with other parasites, host defence mechanisms, and availability of microhabitats [9]. Host selection is an important factor in parasite speciation [38, 64]; for instance, the high motility of many plant-feeding insects allows these parasites to seek out a preferential host species and individuals, while this choice is not available for many other less motile parasite groups that rely on passive transmissions and dispersal pathways. However, the role of host selection as a criterium for sympatric speciation remains a discussion point in the literature [38, 64] as the strict definition of sympatric speciation would potentially limit its applicability to only a few groups of metazoan parasites, such as plant-pathogenic insects [64]. The potentially high extinction rate of parasite species can further complicate inferring their evolutionary history [71, 87]. Consequently, distinguishing between allopatric and sympatric speciation in parasites can be challenging.

### Monogenean flatworms: a model of host-parasite evolution

Monogenean parasites have been suggested as a model system for studying the processes of parasite diversification because of their simple life cycle, morphological and ecological diversity [90], and the high species richness of some genera e.g. *Dactylogyrus* Diesing, 1850 [1, 23], *Gyrodactylus* von Nordmann, 1832 [131, 133], and *Cichlidogyrus* Paperna, 1960 [12, 79]. Monogenean species are often restricted to a few closely related host species [90] or even to microhabitats on a single host species [26,44,63]. African cichlid fishes and the monogeneans belonging to *Cichlidogyrus* are one of most extensively studied fish-monogenean systems and have been proposed as a macroevolutionary model for host-parasite interactions [11,86,92,125]. African cichlids are well-known for their spectacular adaptive radiations [108,114,118], and their role as model for evolutionary research [114]. The cichlid-*Cichlidogyrus* species network is the most extensively described host-parasite network from a species-rich host radiation [10]. A recent meta-analysis counted 477 different host-parasite combinations in this study system [10].

Despite these extensive research efforts, many species of *Cichlidogyrus* remain undiscovered [11, 125]. Most cichlid species have not been examined for parasitic infections, although research spanning several decades has explored species of *Cichlidogyrus* from the East African cichlid radiations (e.g. [27,98,99]) and from the economically relevant tilapias belonging to *Coptodon* Gervais, 1848 and *Oreochromis* Günther, 1889 [21,40,42,77,80,92]. Extensive knowledge gaps remain, especially for species native to Western and Central Africa.

### Chromidotilapiine cichlids: species-rich yet overlooked

Chromidotilapiini Greenwood, 1987 is the most species-rich tribe of cichlids of Central and Western Africa. The tribe includes more species (62) than the tilapias belonging to Oreochromini Dunz & Schliewen, 2010 (59) and Coptodonini Dunz & Schliewen, 2013 (31) across Africa [134]. Nonetheless, the parasite diversity of the latter tribes has been far more extensively studied [11], mainly owing to the economic importance of some of their members [125]. Chromidotilapiines are riverine [111] and also one of the earliest diverging African cichlid lineages together with Tylochromini Poll, 1986, Pelmatochromini Greenwood, 1987, Hemichromini Hoedeman, 1947, and Heterochromidinae Kullander, 1998 [113]. These fishes are the only African cichlids that are not included in the haplotilapiines, a large monophyletic group containing all tilapia-like cichlids and all members of the Eastern African radiations [17].

In terms of their evolutionary history, chromidotilapiines show strong allopatric patterns and their species divergences have been driven by ancient geographic processes rather than ecological specialisation [111]. Many species have somewhat restricted known geographical ranges, e.g., species of *Teleogramma* Boulenger, 1899 as well as *Enigmatochromis lucanusi* Lamboj, 2009, and *Limbochromis robertsi* (Thys van den Oudenaerde & Loiselle, 1971) [52, 111]. Chromidotilapiines are also often geographically separated from their congeners by the limits of river basins (e.g. species of *Benitochromis* Lamboj, 2001*, Congochromis* Stiassny & Schliewen, 2007*, Nanochromis* Pellegrin, 1904*, Pelvicachromis* Thys van den Audenaerde, 1968 [53,55,120], and *Thysochromis emili* Walsh, Lamboj & Stiassny, 2020 [128]). Furthermore, representatives of different genera appear to occupy similar ecological niches in their respective geographical ranges (e.g. the sand-dwellers *Parananochromis longirostris* (Boulenger, 1903) and species of *Nanochromis*; see [111] and references therein for more examples), which seems indicative of allopatric speciation. Phylogenetic analyses highlight that these similar niches in different geographical areas also produced morphologically similar species groups, which have been in the same genera as a consequence. These groups include *Chromidotilapia* sensus stricto in Central Africa and the ‘*Chromidotilapia guntheri* group’ in Western Africa, and *Pelvicachromis* sensu stricto and species of *Pelvicachromis* from the Upper Guinea region [111]. Some species with distribution over large areas have in fact been found to consist of groups of morphologically similar species separated by geographical barriers, e.g. a population of *Pelvicachromis taeniatus* Boulenger, 1901 from Cameroon was found to be morphologically similar to, yet distinct from populations from Benin and Nigeria, therefore, the former was reassigned the name *P. kribensis* Boulenger, 1911 [55]. Hemichromine cichlids, a group that shares much of its geographical range with the chromidotilapiines presents similar cases, where geographically separated groups proved to be distinct species [4].

Of all 78 described species belonging to Chromidotilapiini, parasites have been reported of only three species [*Chromidotilapia guntheri* (Sauvage, 1882), *Parananochromis caudifasciatus* (Boulenger, 1913) and *Benitochromis batesii* (Boulenger, 1901)] [2,15,72]. In the present study, we investigate the morphological evolution of the monogenean fauna of species belonging to the genera *Chromidotilapia*, *Congochromis*, *Divandu*, *Nanochromis*, *Parananochromis*, *Pelvicachromis,* and *Thysochromis.* As these species have not previously been examined for parasites, we expect to find new species as monogeneans express a high level of host specificity [127]. Furthermore, we expect to detect strong allopatric speciation patterns similar to those observed in the host lineages. The present study will expand our knowledge on cichlid-*Cichlidogyrus* interactions and the evolutionary history of *Cichlidogyrus*, one of the most species-rich genera of parasites on the African continent.

## Material and methods

### Parasite collection and morphological examination

Fish specimens were obtained from the ichthyological collection of the Royal Museum for Central Africa (RMCA) (Table 1). The gills of 149 individuals belonging to 27 species of Chromidotilapiini collected from several locations in West and Central Africa (Figure 1) were dissected and subsequently stored in 100% ethanol. The gills were then screened under a stereomicroscope for the presence of monogenean infections. Parasite specimens were mounted on slides with a drop of Hoyer’s medium [37] for morphological identification. Parasite identification and description were conducted using the Leica DM 2500 LED microscope (Leica Microsystems, Wetzlar, Germany) on 400x and 1000x magnification. High-resolution images were taken through the software LasX v3.6.0 (Leica Microsystems, Wetzlar, Germany). Type material was deposited in the invertebrate collection of the Royal Museum for Central Africa (Tervuren, Belgium) (RMCA) (RMCA_VERMES_XXXXX–XXX), the collection of the research group Zoology: Biodiversity and Toxicology of Hasselt University (Diepenbeek, Belgium) (HU XXX–XXX), the Finnish Museum of Natural History (Helsinki, Finland) (MZH XXXXX–XXX), and the Iziko South African Museum (Cape Town, South Africa) (SAMC-XXXXXXX-XXX).

**Figure 1.**
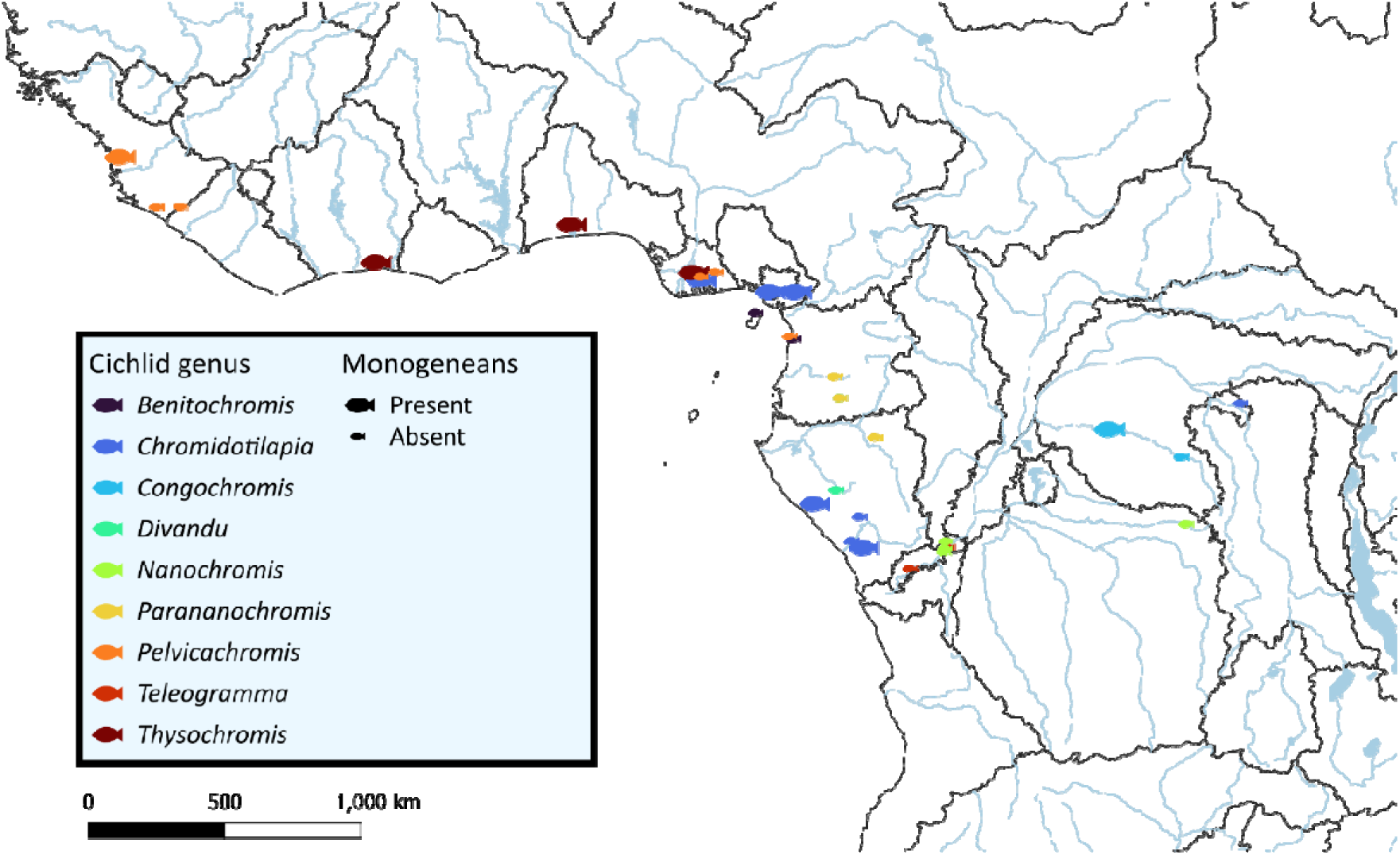
Sampling locations of chromidotilapiine cichlids across Central and West Africa with the presence of monogeneans indicated by size of the symbols. Borders indicate limits freshwater ecoregions according to Thieme et al. [121].

**Table 1.**
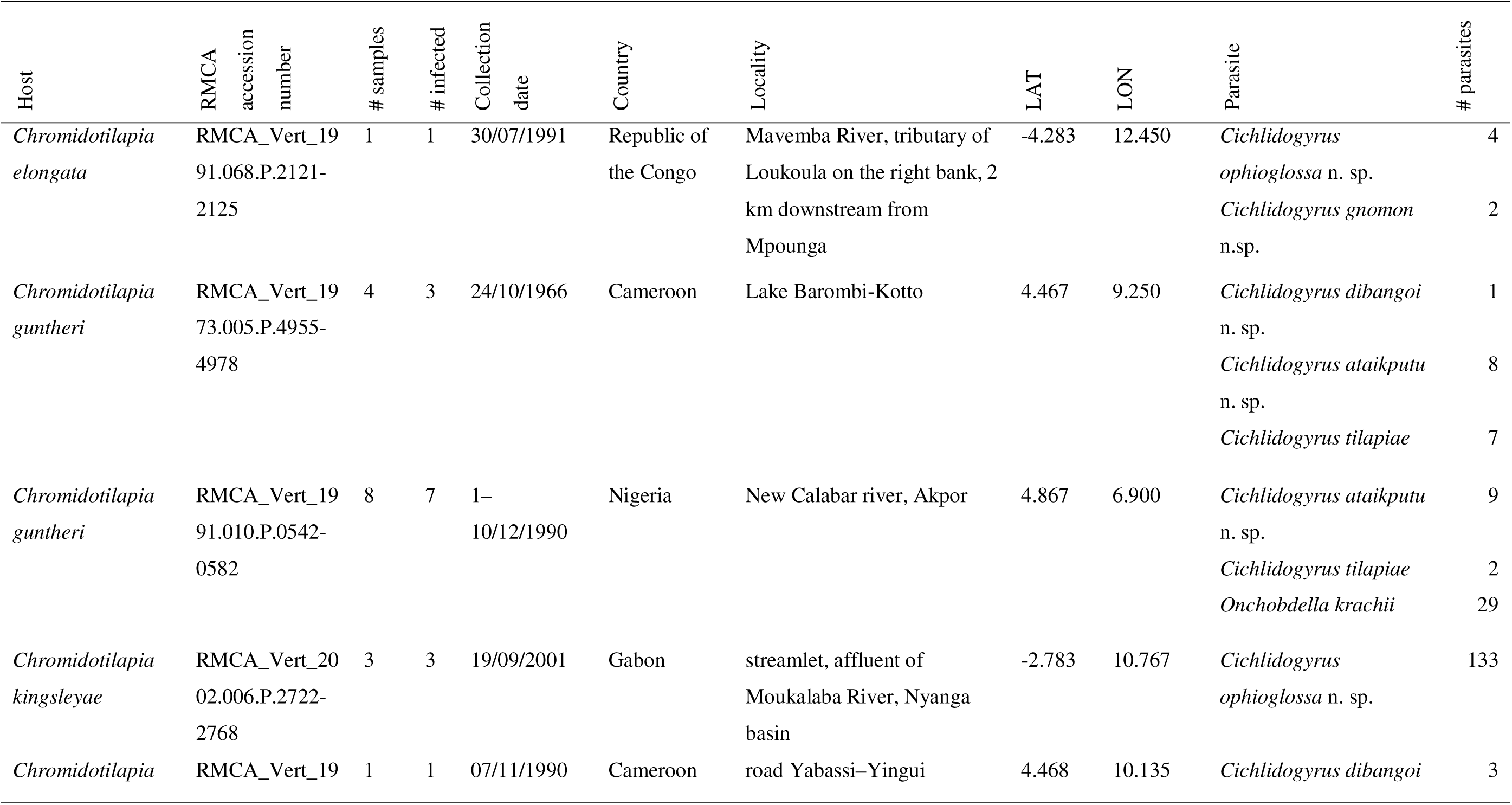
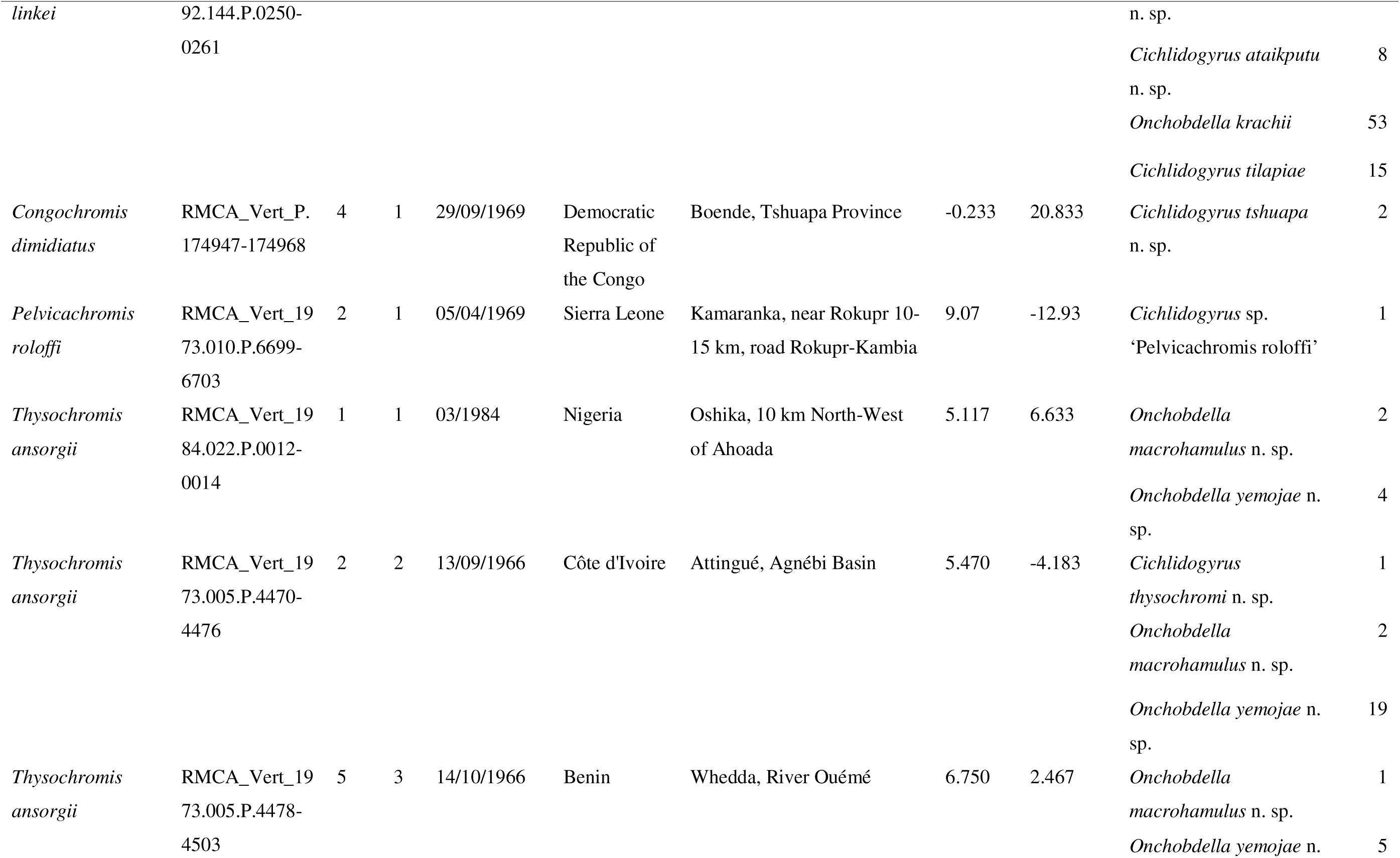
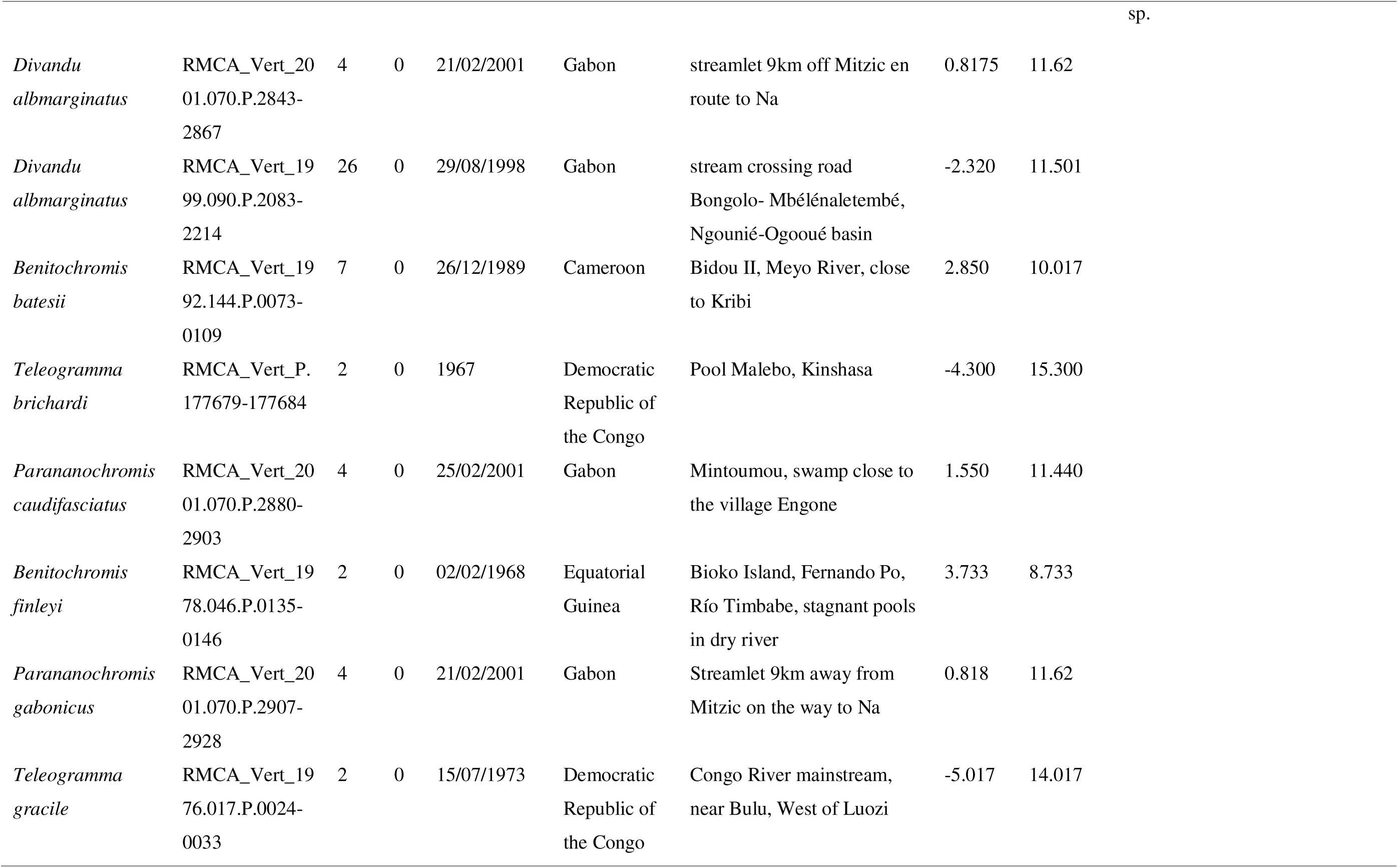
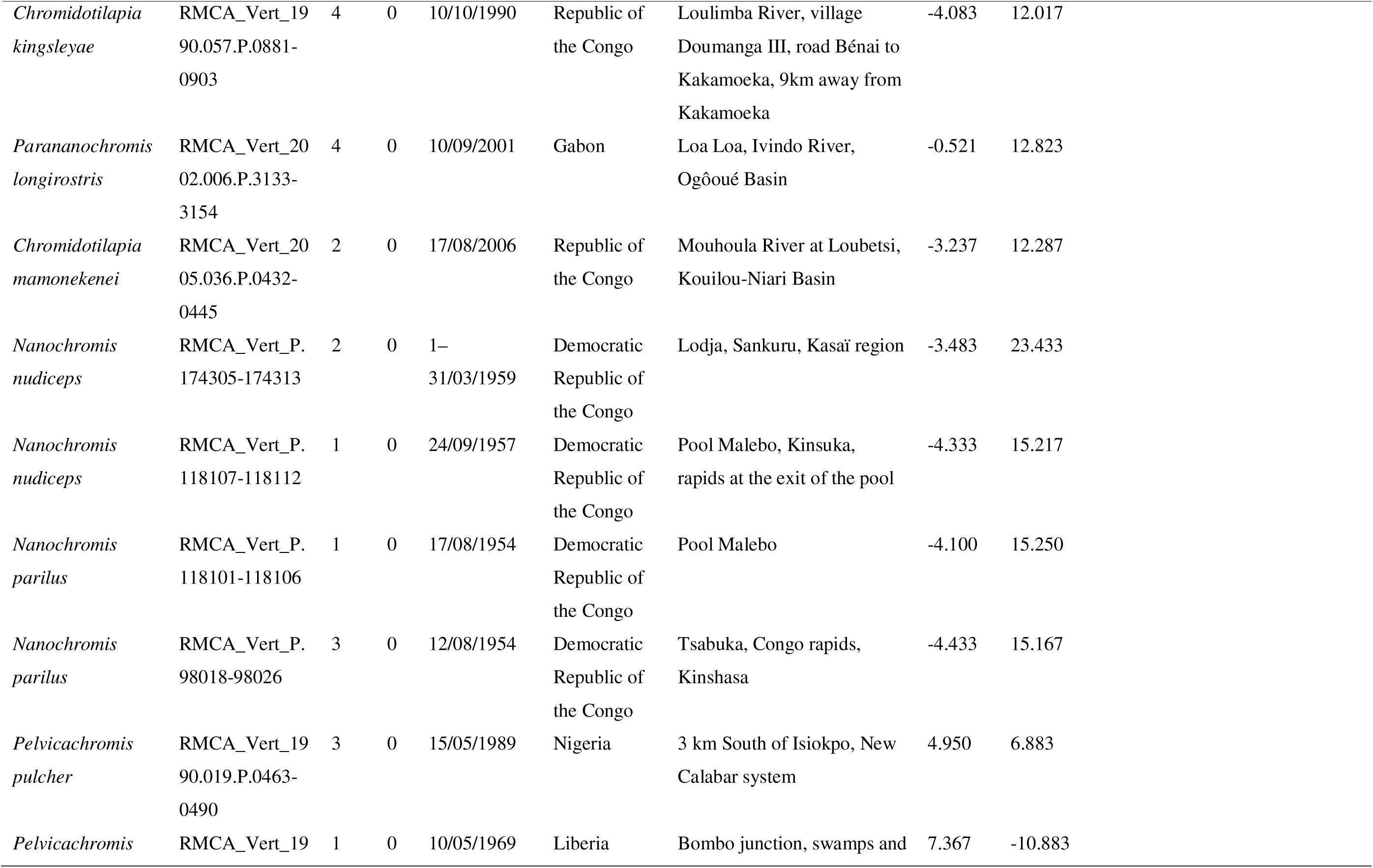
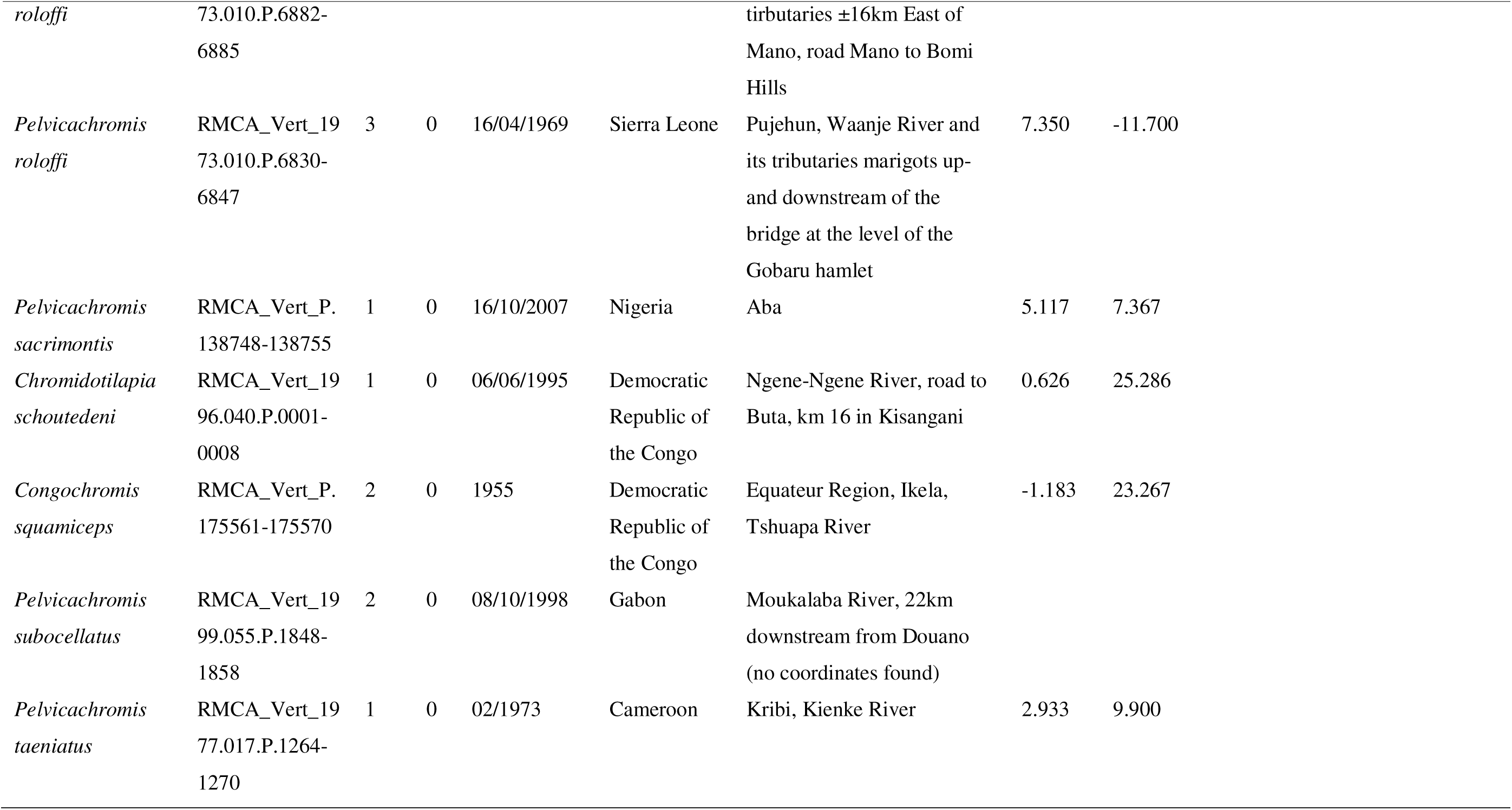
Sampling sites and dates of host specimens in the collection of the Royal Museum for Central Africa (RMCA) in Tervuren, Belgium, and monogenean gill parasites found on their gills.

### Morphometrics, missing data, and principal component analysis

Species characterisations of dactylogyrid monogenean species are frequently based on the morphology of the sclerotised structures of the attachment and reproductive organs [96]. Therefore, parasites were grouped according to phenotypic characters in these structures. For an analysis of morphometric characters, we also took 29 different measurements of the hard parts of the haptor, the male copulatory organ (MCO) and the vagina (Figure 2). The terminology was based on Pariselle et al. [78]. The marginal hooks are counted according to Llewellyn [59].

**Figure 2:**
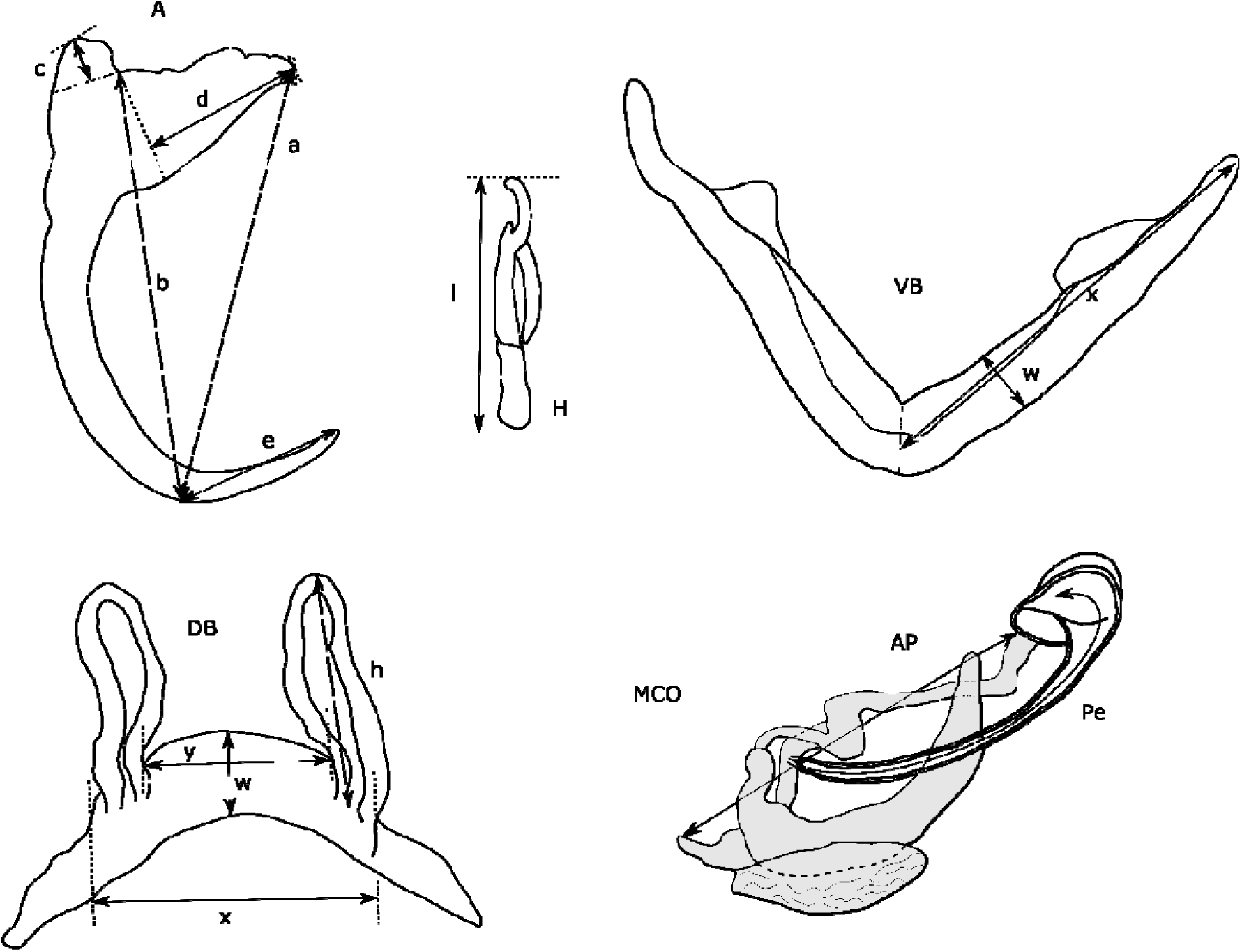
Measurements used for sclerotised structures of haptor and reproductive organs of *Cichlidogyrus* sp. VA, ventral anchor, DA, dorsal anchor: a, Total length, b, Length to notch, c, Outer root length, d, Inner root length, e, Point length; H, Hook length; VB Ventral bar: x Branch length, w, Branch width; DB Dorsal bar: h, auricle length, w, maximum straight width, x, total length, y, distance between auricles; MCO Male copulatory organ: AP, accessory piece straight length, Pe, copulatory tube curved length.

The monogeneans were identified to species level based on shapes and sizes of the sclerotised structures. However, to assess additional morphological variation in similar species, we conducted principal component analyses (PCA). Standard PCA approaches require a dataset without missing information but often measurements of monogenean flatworms are incomplete due to the fragility of the worms and damage caused to the samples by the mounting process. Multiple methods have been proposed to address missing data in PCAs (see [119]). Here, we employ non-iterative partial least squares (NIPALS) with Gram–Schmidt orthogonalization as implemented in the *R* package *nipals* v0.8 [130].

### Phylogenetic position: Maximum parsimony and machine learning

We used the morphometric data to infer the phylogenetic position of the new species of *Cichlidogyrus* based on the dataset published by Cruz-Laufer et al. [12]. Cruz-Laufer et al. [12] demonstrated that morphometric data of the attachment and reproductive organs can indicate phylogenetic relationships between species of *Cichlidogyrus*, albeit limited to certain measurements and groups of related species. Here, these data were reanalysed by expanding the parsimony and machine learning approaches to the new species found on chromidotilapiine cichlids.

First, phylogenetic positions were inferred based on the morphometric measurements (Figure 2) and the morphological discrete characters for the reproductive organs suggested by Cruz-Laufer et al. [12] (Table 2). In some cases, we proposed new character states for the new species (Table 2). Phylogenetic inference was performed under maximum parsimony in TNT v1.5 [32, 33] and with the latest genus-wide molecular phylogeny [12] used as a backbone (options *force* and *constrain*) to place the new species amongst their congeners with published DNA sequences. We applied extended implied weighting (option *xpiwe*) to reduce the impact of missing data [28] that were weighted artificially high in the original implied weighting method [29]. Furthermore, tree topologies were inferred for a range of values for the concavity constant k (20, 21, 23, 26, 30, 35, 41, 48, 56) to infer the most stable tree topology. We assigned each character a separate weight as recommended for continuous data [34]. As suggested by Mirande [68], we selected values of k that resulted in the highest distortion coefficient and subtree pruning and regrafting (SPR) distance on average compared to the other consensus trees. The final consensus tree was inferred from trees produced under the optimised k values. Tree searches involved rounds of tree fusing, sectorial searches, and tree drifting [30] under default settings with each round stopped after three hits of the same optimum. Gaps were treated as missing data. Branch support was estimated through symmetric resampling (probability of change: 0.33) and values expressed as differences in frequencies (GC: ‘Groups present/Contradicted’) as implied weighting methods can distort bootstrapping and jackknifing methods [31].

**Table 2.** List of character states of reproductive organs used for parsimony analysis. A new character was used to capture the unique wing-like structure associated with the accessory pieces of Cichlidogyrus ophioglossa n. sp. and C. gnomon n. sp. (in bold: shape of accessory piece—o).

| Character | Character states |
| --- | --- |
| Shape of copulatory tube | a. Straight: penis more or less straight with no strong arching, twisting, looping, or spiralling but can be slightly sinuous or arched. |
|  | b. Straight, thick-walled: same as before but wall of penis present thickening. |
|  | c. Arched: penis strongly arched in one direction, distal portion often held in position by accessory piece. |
|  | d. Looped: penis draws a loop in the shape of a G. |
|  | e. Large loop: penis draws large circle ending in distal portion of accessory piece. |
|  | f. Spiralled: penis draws spiral in large radius. |
|  | g. Spirally coiled: penis draws spiral in small radius in the shape of a helix |
| Diameter of copulatory tube | a. Tubular: penis in the shape of a simple tube. |
|  | b. Widened: penis widened. |
|  | c. Bulbous: penis presents a bulbous portion (outside the basal bulb). |
| Shape of accessory piece | a. Simple: elongated accessory piece without additional structures mentioned in the other character states but species with more unique structures such as connecting stalks and caps are also included here. |
|  | b. Furcated: accessory piece present one or more furcations. |
|  | c. Distal hook: Accessory piece ends in a single distal hook. |
|  | d. Distal flap: Accessory piece ends in a single distal flap. |
|  | e. Gutter-like: Accessory piece in the shape of a gutter guiding the penis. |
|  | f. Ribbon-like: Accessory piece is a flattened structure in the shape of a ribbon or drape. |
|  | g. Spirally coiled: Accessory piece in the shape of simple helix. |
|  | h. Looped: Accessory piece draws a loop in the shape of a G. |
|  | i. Reduced: Accessory piece reduced to a thin, string-like structure or absent. |
|  | j. Complex, S-shaped: massive, roughly S-shaped accessory piece (Fig. 5: *CPO*) that is frequently connected to the heel. The accessory piece has an extension or thickening at the first turn in the proximal half and frequently displays a folded back, straight and pointy, or hook-like distal end, or sometimes additional terminations resulting in a furcate ending with two or three digitations. However, the first turn is never V-shaped or knee-like such as in (l) and the hook-shaped termination is never sickle-like such as in (c). |
|  | k. Complex, C-shaped: complicated roughly C-shaped accessory piece often |
|  | with finger or hook-shaped outgrowths and marked heel (Fig. 5: *OreoI*). |
|  | l. Two portions, V-shaped: accessory piece consists of two distinct portions shaped like a V with an expanded knee-like bend (Fig. 5: *Hemi*) |
|  | m. Two portions, spiralling: accessory piece consists of two distinct portions, large spiral followed by non-spiralled distal portion. |
|  | n. In two parts: accessory piece consists of two distinct, superimposed parts. |
|  | **o. Complex with associated wing-like structure: accessory piece with multiple processes and associated wing-like structure** |
| Shape of vagina | a. Non-sclerotised: Vagina not sclerotised. |
|  | b. Tubular: Vagina in the shape of a simple tube. |
|  | c. Bulbous: Vagina widened in at least one portion. |
|  | d. Spiralled: tubular vagina that draws a spiral. |

Second, species of *Cichlidogyrus* characterised here were placed in groups of related species of *Cichlidogyrus* reported by Cruz-Laufer et al. [12] using supervised machine learning (ML). Machine learning algorithms improve prediction accuracy through experience, i.e. repetition. Here, we trained ML algorithms to classify specimens in species groups based on their morphology. Cruz-Laufer et al. [12] reported a moderate performance of ML algorithms. However, their study only included one type of ML algorithm (support vector machines) and their algorithm was trained only on all available continuous morphometrics. In contrast, we applied three widely used ML algorithms including random forest (RF), support vector machines (SVM) with radial basis kernel function, and artificial neural networks (ANN) to all morphometric measurements combined as well as the discrete morphological characters of the reproductive organs proposed by Cruz-Laufer et al. [12]. This analysis was conducted in the *R* package *caret* (Kuhn, 2008) using the methods *rf* [57], *svmRadial* [67], and *nnet* [126]. Missing data were imputed through k-nearest neighbour imputation, centred, and scaled through the function *preProcess*. Tuning parameters were optimised through grid search (Table 3) tenfold cross-validation with ten repetitions. Model performance was assessed through Cohen’s κ to account for the class imbalance in the data [56]. Following Landis and Koch [56], we considered κ < 0.2 a *slight*, κ between 0.2 and 0.4 a *fair*, κ between 0.4 and 0.6 a *moderate*, κ between 0.6 and 0.8 a *substantial*, and κ > 0.8 an *almost perfect* agreement.

**Table 3.** Overview of range of values used for parameter tuning through grid search for different machine learning algorithms

| Algorithm | Parameters | Values for grid search |
| --- | --- | --- |
| Support vector machines | C | $2^{-15}, 2^{-13}, 2^{-11}, 2^{-9}, 2^{-7}, 2^{-5}, 2^{-3}, 2^{-1}, 2^1, 2^3$ |
| | $\sigma$ | $2^{-5}, 2^{-3}, 2^{-1}, 2^1, 2^3, 2^5, 2^7, 2^9, 2^{11}, 2^{13}, 2^{15}$ |
| Artificial neural networks | size | 3, 5, 10, 20 |
|  | decay | 0.5, 0.1, 1E-2, 1E-3, 1E-4, 1E-5, 1E-6, 1E-7 |
| Random Forest |  | 1, 2, 3, 4, 5, 6, 7, 8, 9, 10, 11, 12, 13, 14, 15, 16, |
|  | mtry | 17, 18, 19, 20, 21, 22, 23, 24, 25 |

## Results

### Morphological examination

Of the 27 fish species examined, specimens of nine species were infected with monogenean flatworms, including *Chromidotilapia elongata* Lamboj, 1999, *Chromidotilapia guntheri* (Sauvage, 1882), *Chromidotilapia guntheri loennbergii* (Trewavas, 1962), *Chromidotilapia kingsleyae* Boulenger, 1898, *Chromidotilapia linkei* Staeck, 1980, *Congochromis dimidiatus* (Pellegrin, 1900)*, Pelvicachromis roloffi* (Thys van den Audenaerde, 1968) and *Tysochromis ansorgii* (Boulenger, 1901). We found a total of 6, 69, 45, 135, 118, 2, 2, and 65 monogenean parasites respectively. Seven species were found to be new to science, of which five belonging to *Cichlidogyrus* and two belonging to *Onchobdella*. Specimens of *Cichlidogyrus tilapiae* (Paperna, 1960) [73] and *Onchobdella krachii* Paperna, 1968 [72] were also found. Species descriptions and characterisations are presented in the following. Note that the authors of the new species are different from the authors of the article [according to Article 50 of the International Code of Zoological Nomenclature (ICZN)]. The Life Science Identifier (LSID) of the article is urn:lsid:zoobank.org:pub:XXXXXXXX. The LSIDs of the new species are reported below. Infection parameters can be found in Table 1. Measurements of species of *Cichlidogyrus* and *Onchobdella* can be found in Table 4 and Table 5 respectively. Symbiotypes and symbioparatypes are given as following: RMCA accession number (specimen IDs).

**Table 4.**
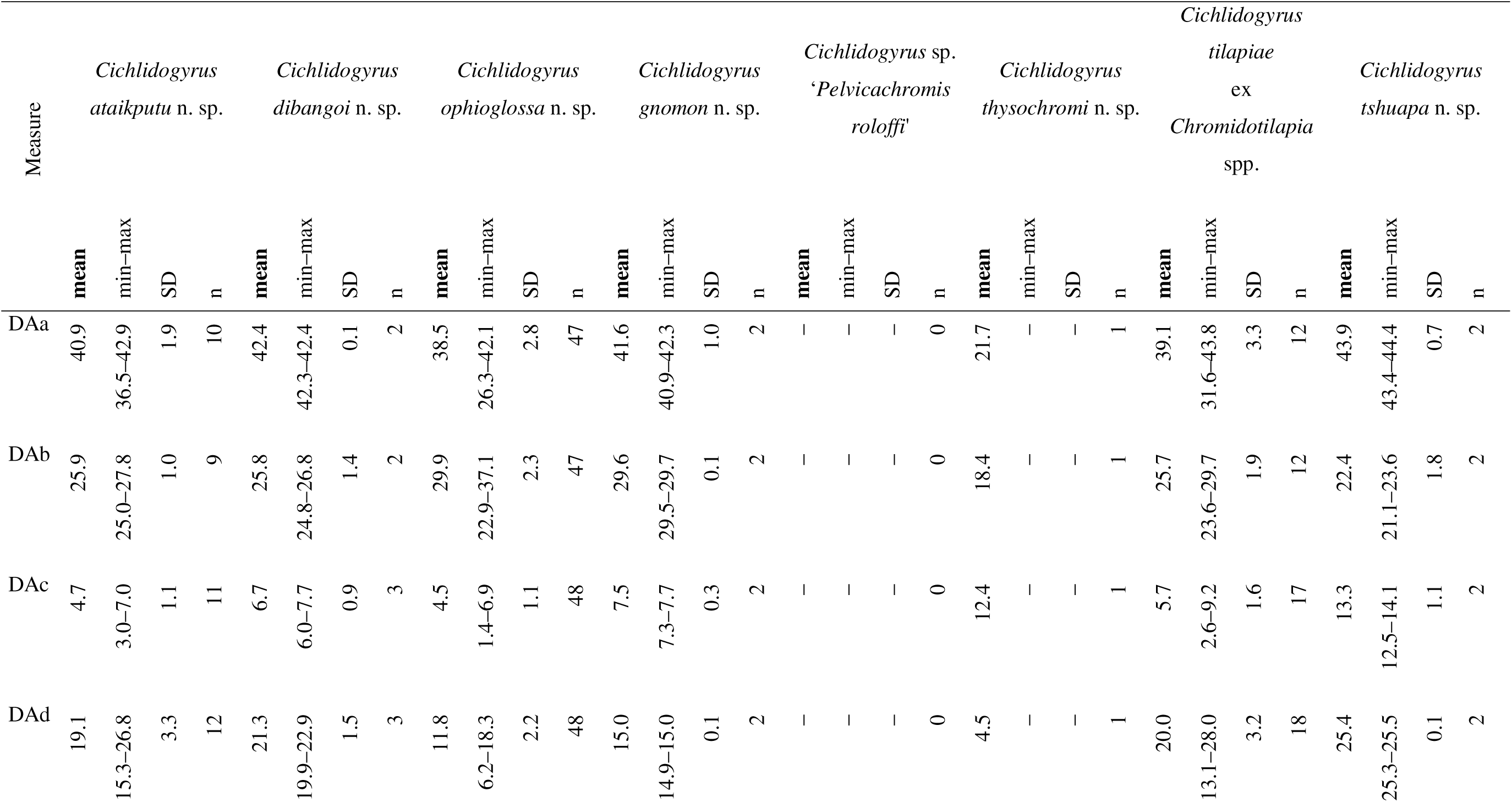
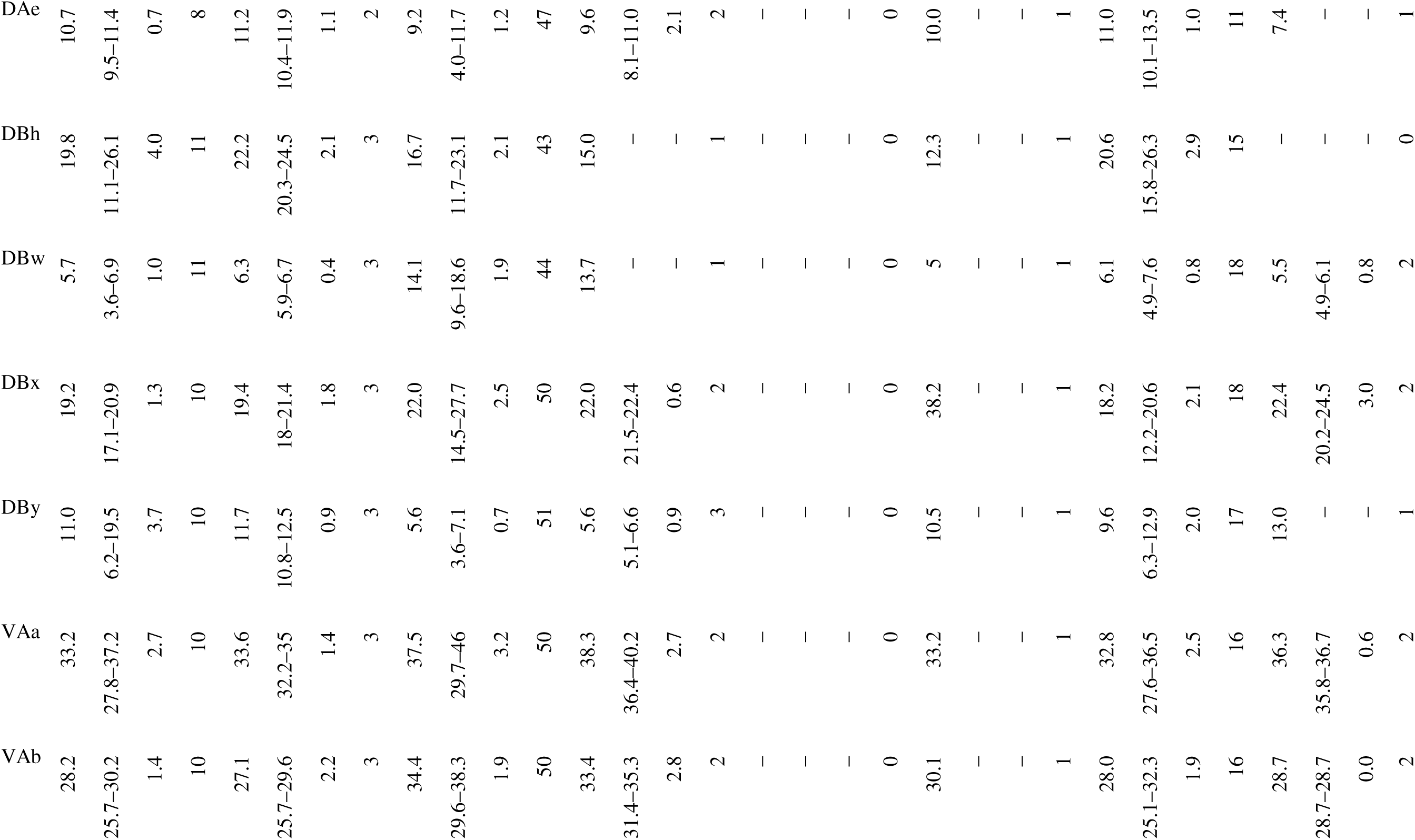
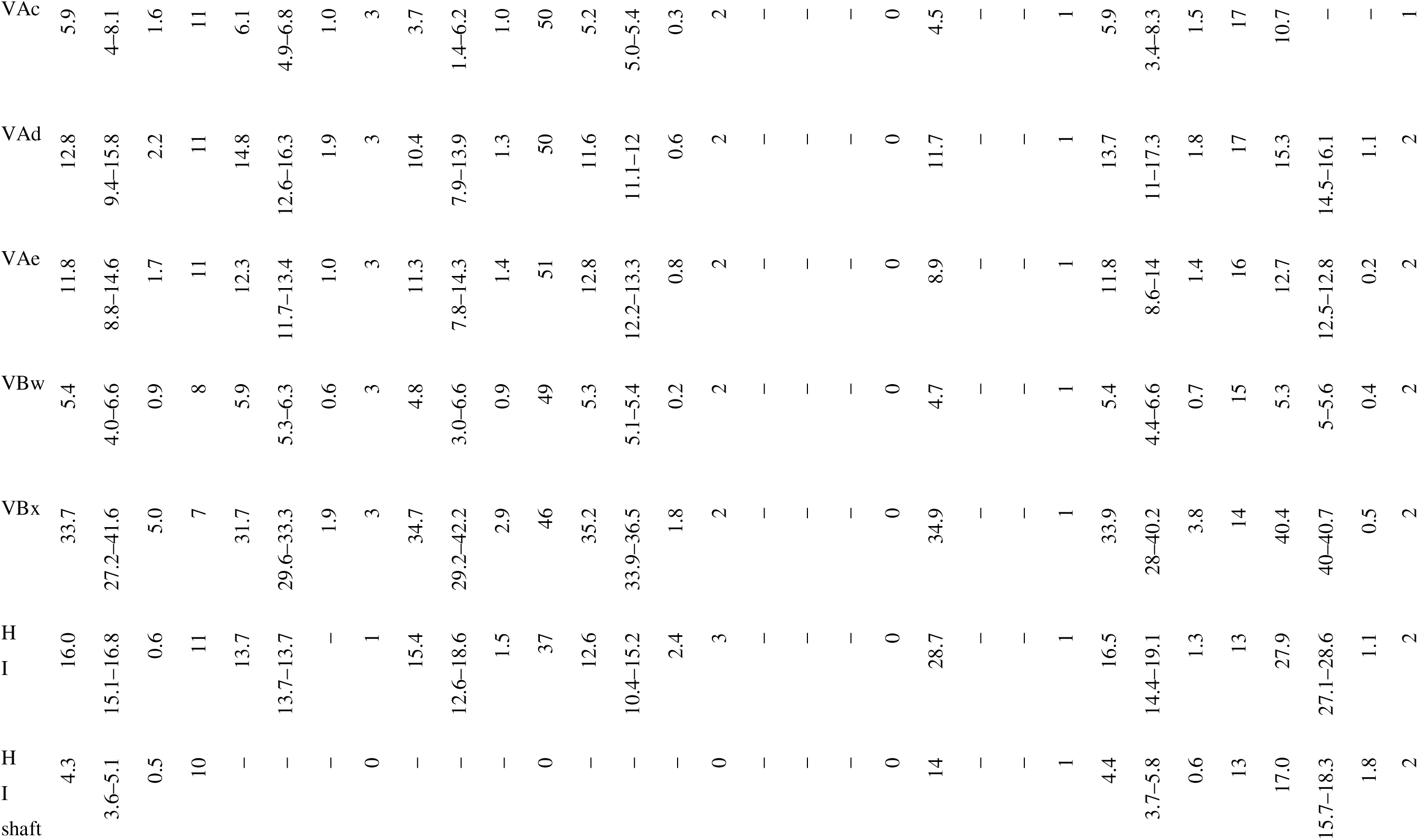
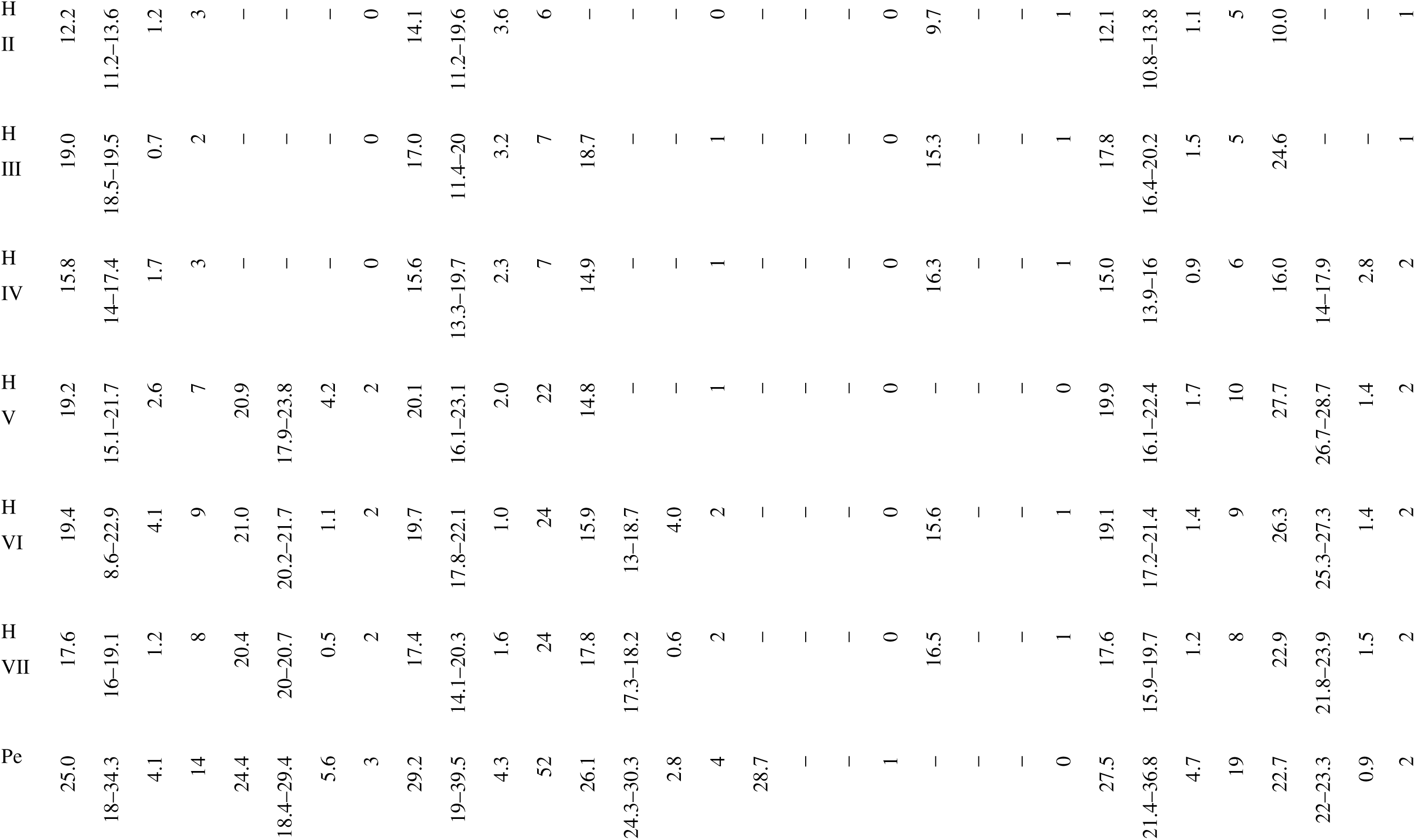
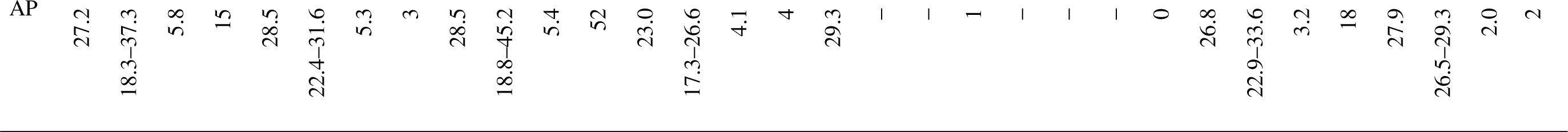
Morphometrics of species of Cichlidogyrus infecting chromidotilapiines cichlids. min–max, minimum and maximum value; SD, standard deviation; n, sample size.

**Table 5.** Morphometrics of species of *Onchobdella* infecting chromidotilapiines cichlids. min–max, minimum and maximum value; SD, standard deviation; n, sample size.

| Measure | *Onchobdella krachii* |  |  |  | *Onchobdella macrohamulus* n. sp. |  |  |  | *Ochobdella yemojae* n. sp. |  |  |  |
| --- | --- | --- | --- | --- | --- | --- | --- | --- | --- | --- | --- | --- |
|  | mean | min–max | SD | n | mean | min–max | SD | n | mean | min–max | SD | n |
| DAa | 37.6 | 29.1–46.8 | 4.6 | 51 | 120.0 | 95.6–131.7 | 14.1 | 5 | 39.4 | 33.2–45 | 3.6 | 20 |
| DAb | 26.7 | 19.4–31.1 | 2.6 | 49 | 102.5 | 92–107.2 | 6.2 | 5 | 29.5 | 24.3–34.6 | 3 | 19 |
| DAc | 2.8 | 1.1–5.1 | 0.9 | 47 | 4.3 | 2.3–7 | 1.8 | 5 | 2.2 | 1–5.5 | 1.4 | 18 |
| DAd | 15.9 | 10.2–22.2 | 2.8 | 51 | 28.8 | 14.6–45.4 | 13.5 | 5 | 13.7 | 11.3–17 | 1.7 | 19 |
| DAe | 16.3 | 10.8–19.8 | 2.5 | 52 | 28.1 | 24.9–31.2 | 2.6 | 5 | 20 | 15.7–25.2 | 2.4 | 21 |
| DBw | 5.5 | 3.2–7.7 | 1.1 | 57 | 17.8 | 14.2–21.6 | 3.4 | 5 | 5 | 2.9–9.7 | 1.5 | 22 |
| DBx | 41.3 | 23.9–54.6 | 7.3 | 56 | 70.4 | 60.1–79.1 | 7.3 | 5 | 38.5 | 31.4–49.8 | 5.5 | 22 |
| VAa | 13.8 | 11–17.3 | 1.1 | 53 | 15.5 | 14.5–16.4 | 0.9 | 5 | 8 | 6.6–9.7 | 1 | 21 |
| VAb | 11.8 | 9.5–15.1 | 0.8 | 53 | 16.1 | 14.6–18.6 | 1.7 | 5 | 7.9 | 6.6–9.5 | 0.9 | 21 |
| VAc | 2.3 | 1.2–4 | 0.6 | 53 | 3 | 2.5–4.1 | 0.7 | 5 | 1.6 | 0.8–2.3 | 0.4 | 20 |
| VAd | 6.5 | 5.2–8.3 | 0.8 | 53 | 5.3 | 4–8.2 | 1.6 | 5 | 4.1 | 3.3–5.4 | 0.5 | 21 |
| VAe | 6.8 | 5.2–8.8 | 0.8 | 53 | 7.1 | 4.5–9.8 | 2.3 | 5 | 4.2 | 2.6–5.5 | 0.8 | 20 |
| VBw | 3.1 | 1.9–4.8 | 0.6 | 52 | 2.7 | 1.9–3.4 | 1.1 | 2 | 1.5 | 1.3–1.8 | 0.2 | 11 |
| VBx | 30.5 | 19.2–41.6 | 4.6 | 52 | 42.5 | 40.4–44.6 | 3 | 2 | 20.2 | 16.3–22.4 | 1.8 | 9 |
| H | 14.3 | 11.9–16.3 | 1.2 | 19 | 16.5 | 15.7–16.9 | 0.5 | 4 | 13.4 | 10.5–15.5 | 1.9 | 5 |
| Pe | 53.3 | 33.1–99.8 | 11.8 | 66 | 65.4 | – | 1 | 2 | 36.6 | 28.8–45.2 | 4.7 | 17 |
| AP | 21.1 | 14.1–41.6 | 3.8 | 63 | 32.3 | 27.7–36.8 | 6.4 | 2 | 30.3 | 21–44.8 | 6.8 | 17 |

Subclass Polyonchoinea Bychowsky, 1937

Order Dactylogyridea Bychowsky, 1937

Family Dactylogyridae Bychowski, 1933

Genus Cichlidogyrus Paperna, 1960

**Name**: *Cichlidogyrus ataikputu* Moons, Kmentová, Pariselle, Vanhove & Cruz-Laufer n. sp.

**Type-host**: Chromidotilapia guntheri (Sauvage, 1882)

**Additional host***: Chromidotilapia linkei* Staeck, 1980

**Locality**: New Calabar river, Akpor, Nigeria; 4.87, 6.9; 01/12/1990 on type host

**Additional locality**: Lake Barombi-Kotto, Cameroon on *Chromidotilapia guntheri* and road Yabassi-Yingui, Cameroon on *Chromidotilapia linkei*

**Symbiotype**: RMCA_Vert_1991.010.P.0542-0582 (578)

**Symbioparatype**: RMCA_Vert_1991.010.P.0542-0582 (576, 577, 581, 582); RMCA_Vert_1973.005.P.4955-4978 (CGL9, CGL16,CGL20); RMCA_Vert_1992.144.P.0250-0261 (B, C)

**Type-material**: Holotype—RMCA_VERMES_XXXXX, Paratypes—RMCA_VERMES_XXXXX–XXX, HU XXX–XXX, MZH XXXXX–XXX, SAMC XXXXXXX–XXX

**ZooBank registration:** Life Science Identifier (LSID) for *Cichlidogyrus ataikputu* Moons, Vanhove, Pariselle & Cruz-Laufer n. sp.—urn:lsid:zoobank.org:act:XXXXXXXXXXXX.

**Site in host**: Gills

**Etymology**: The species epithet ‘*ataikputu*’ correctly spelled ‘ata ikputu’ is Igbo, a language spoken in the area where the holotype has been sampled. ‘ata’ translates to ‘consumes’, whereas ‘ikputu’ refers to Gunther’s mouthbrooder (*Chromidotilapia guntheri*) [123].

**Description:** [based on 15 specimens, Figure 3] Two pairs of anchors. Ventral anchor with a more developed inner root than outer root and deep indentation. Dorsal anchors with well-developed inner root. Sturdy ventral transverse bar V-shaped with small membranous attachment at the base of the branches. Dorsal transverse bar with thick midsection and long and slender auricles. Marginal hooks seven pairs, all approximately the same size except for pair 2, which is smaller. Male copulatory organ (MCO) consists of a copulatory tube and accessory piece. The copulatory tube has a broad base and becomes slender and curved towards the distal end, where it is guided by a sheath-like portion the accessory piece. The accessory piece is as broad as the copulatory tube and is attached to the base of the copulatory tube. At the distal end, it bends at a 90° angle and follows the copulatory tube terminating in two small pointy protuberances. The accessory piece folds halfway and guides the copulatory tube. No heel present. No sclerotised vagina observed.

**Figure 3:**
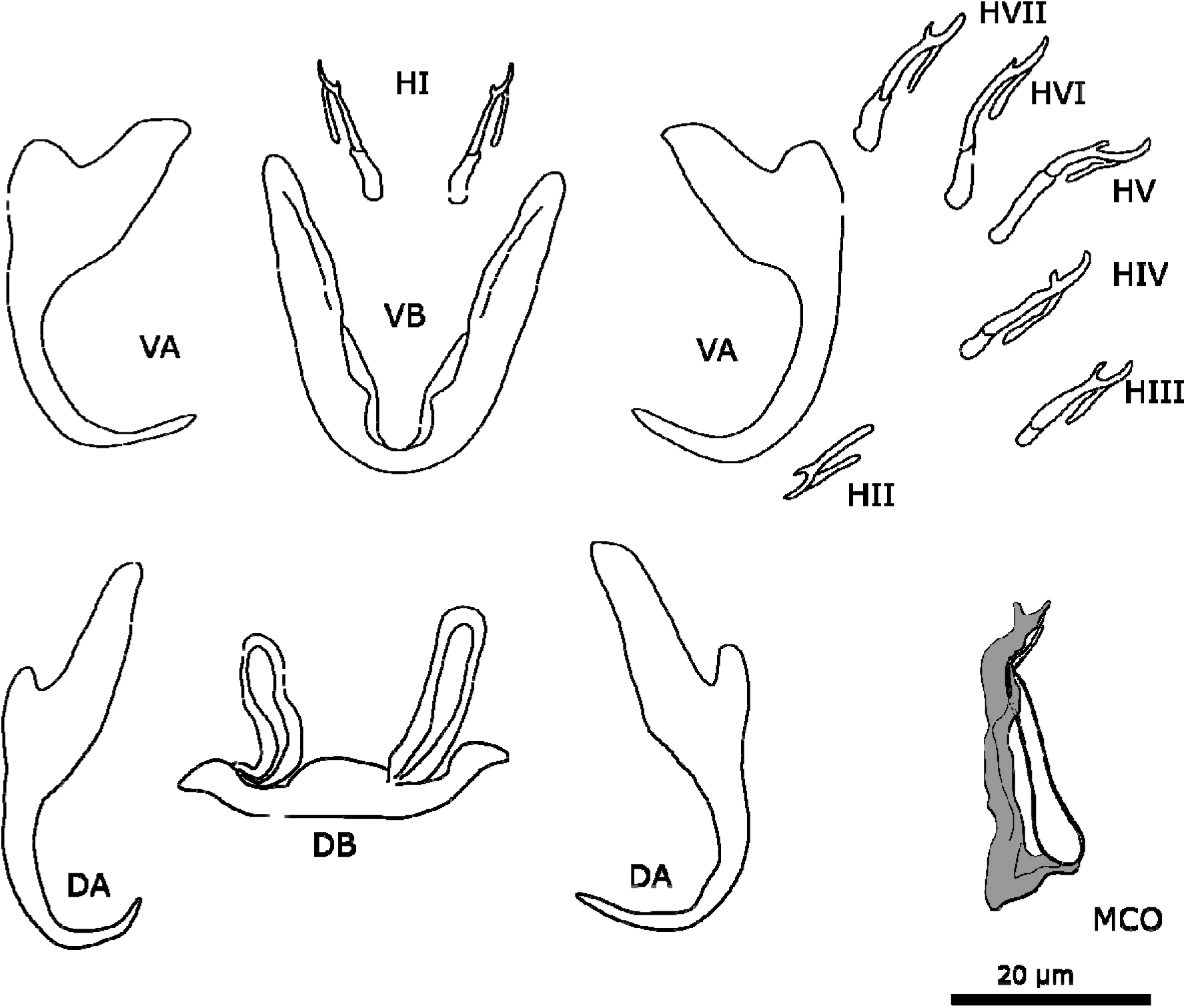
Sclerotised structures of *Cichlidogyrus ataikputu* n. sp. Abbreviations: HI-HVII, hooks; VA, ventral anchor; VB, ventral transverse bar; DA, dorsal anchor; DB, dorsal transverse bar; MCO, male copulatory organ.

**Remarks:** The specimens show typical features of species of *Cichlidogyrus*, e.g. (i) two pairs of anchors (one ventral and one dorsal), two transverse bars (V-shaped ventral bar, doral bar with two auricles); (ii) seven pairs of marginal hooks; (iii) a MCO consisting of a copulatory tube and generally an accessory piece; and (iv) a vagina, which can be sclerotised [73, 79]. *Cichlidogyrus ataikputu* n. sp. presents similarities with *Cichlidogyrus tilapiae*. The dorsal anchors of the two species are similar in having a well-developed inner root and a reduced outer root. The dorsal bars are also similarly shaped, as are the lengths of the auricles of in *C*. *ataikputu* n. sp. (20.3–24.5 µ m) and *C. tilapiae* (23–34 µ m) according to Rindoria et al. [102] (but not the original measurements by Paperna [73]: 9–19 µ m). The differences in sizes might be explained by an adaptation to different host species or geographical variation, being *Oreochromis niloticus* (Linnaeus, 1758) and *Sarotherodon galilaeus* (Linneaus, 1758) in Dor, Israel [73]; and *O. leucostictus* (Trewavas, 1933) and *O. niloticus* Lake Naivasha, Kenya [102]. *Cichlidogyrus ataikputu* n. sp. also resembles *Cichlidogyrus dibangoi* n. sp., also described in the present study (see below). The auricles are longer in *C*. *dibangoi* n. sp. The hooks of *C. dibangoi* n. sp. and *C. ataikputu* n. sp. are very similar in morphology and size. At the distal end, the accessory piece of *C. dibangoi* n. sp. encompasses the copulatory tube like a sheath from one side. This sheath-like portion of the accessory piece is also seen in *C*. *ataikputu* n. sp. but is shorter than in *C. dibangoi* n. sp. The end of the accessory piece shows two small protuberances whereas in *C. dibangoi* n. sp., the end is hook-shaped. The morphology of the copulatory tube is similar in C. *dibangoi* n. sp. and *C*. *ataikputu* n. sp., in having a bulbous base followed by a long slender tube. Yet the tube curves at the distal end in *C*. *ataikputu* n. sp., and no heel is present unlike in C. *dibangoi* n. sp.

**Name**: *Cichlidogyrus dibangoi* Moons, Kmentová, Pariselle, Vanhove & Cruz-Laufer n. sp.

**Type-host**: *Chromidotilapia guntheri* (Sauvage, 1882)

**Additional host**: *Chromidotilapia linkei* Staeck, 1980

**Locality**: Lake Barombi-Kotto, Cameroon; 4.47, 9.25; 24/10/1966

**Additional locality**: road from Yabassi to Yingui, Cameroon; on *Chromidotilapia linkei*

**Symbiotype**: RMCA_Vert_1973.005. P.4955-4978 (CGL16)

**Symbioparaype**: RMCA_Vert_1992.144.P.0250-0261 (B)

**Type-material**: Holotype—RMCA_VERMES_XXXXX, Paratypes—RMCA_VERMES_XXXXX, HU XXX

**ZooBank registration:** Life Science Identifier (LSID) for *Cichlidogyrus dibangoi* Moons, Vanhove, Pariselle & Cruz-Laufer n. sp.—urn:lsid:zoobank.org:act:XXXXXXXXXXXX.

**Site in host:** Gills

**Etymology**: The species epithet ‘*dibangoi*’ honours Manu Dibango, a famous saxophonist and singer-songwriter from Cameroon, who incorporated Jazz and traditional Cameroonian elements into his music.

**Description:** [based on 3 specimens, Figure 4] Two pairs of anchors. Ventral anchors with a more developed inner root than outer root. Dorsal anchors have a well-developed inner root and outer root about the same size as the outer root of the ventral anchor. Sturdy ventral transverse bar V-shaped with membranous attachment towards the distal end of the branches. Dorsal transverse bar with a thick middle section and elongated slender auricles. Seven pairs of marginal hooks have approximately the same size, except for pair 2 which is smaller; measurements of pair 3 and 4 could not be assigned due to distortions of the material during the mount process. The MCO consists of a copulatory tube, an accessory piece, and a small heel. The copulatory tube is broad at the base, narrows towards the distal end with a terminal opening. The accessory piece is attached to the base of the copulatory tube. The proximal part of the accessory piece folds towards the copulatory tube. Distally, the accessory piece widens, then narrows again towards a hook-shaped distal end. The accessory piece folds at the mid-portion of the copulatory tube. No sclerotised vagina observed.

**Figure 4:**
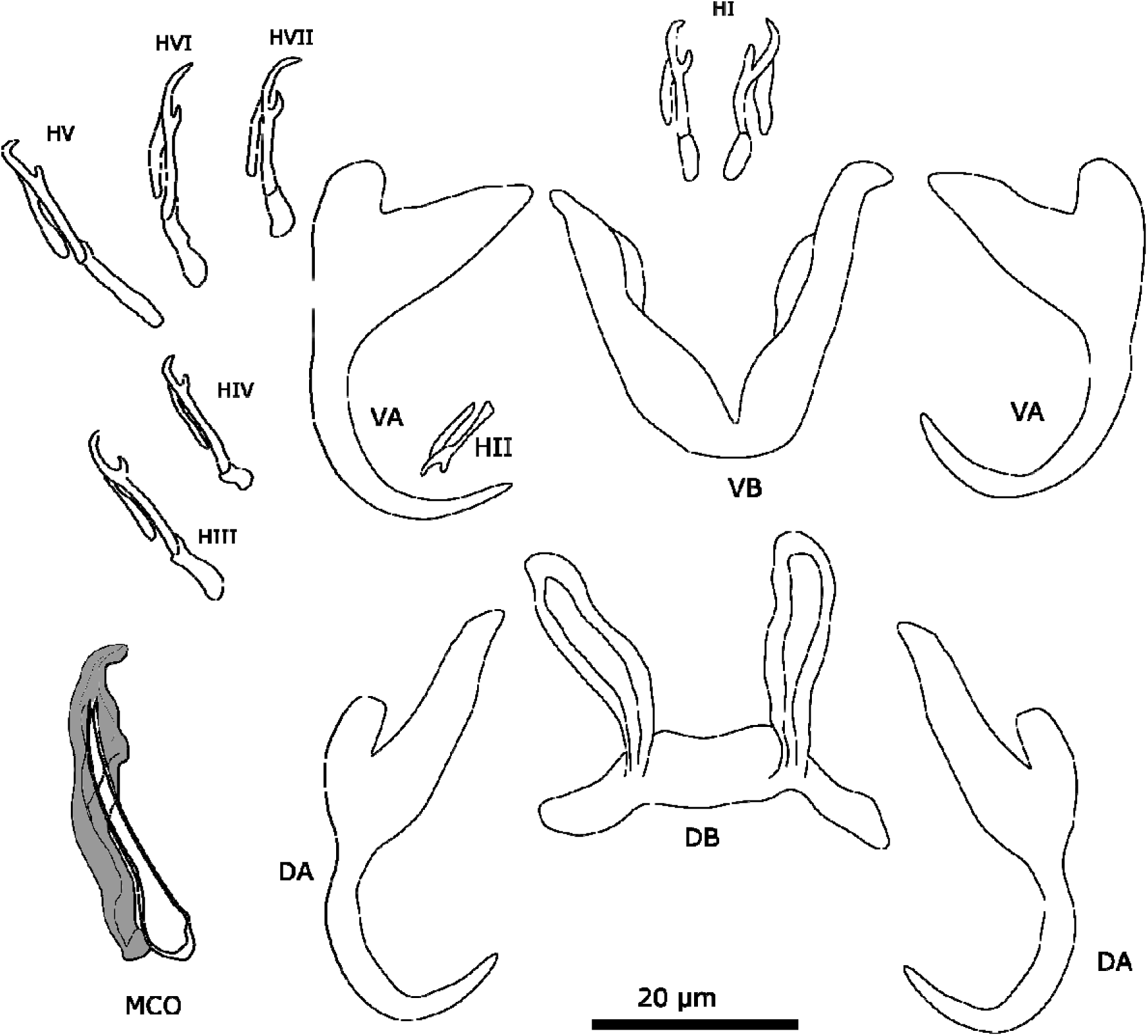
Sclerotised structures of *Cichlidogyrus dibangoi* n. sp. Abbreviations: HI-HVII, hooks; VA, ventral anchor; VB, ventral transverse bar; DA, dorsal anchor; DB, dorsal transverse bar; MCO, male copulatory organ.

**Remarks**: All specimens show diagnostic features of species of *Cichlidogyrus* (see “Remarks” *C. ataikputu* n. sp.). *Cichlidogyrus dibangoi* n. sp. resembles *C. tilapiae*, which infects a wide array of cichlid and non-cichlid hosts [12], and *Cichlidogyrus ataikputu* n. sp. The ventral anchors are morphologically similar to *C. tilapiae* and *C. ataikputu* n. sp. in their size and lengths of their roots. Furthermore, the three species have a dorsal bar that is similar in size with long slender auricles. The auricles are slightly longer in *C. dibangoi* n. sp (20.3–24.5 µ m), than in *C. tilapiae* (9–19 µ m) described by Paperna [73], but not longer than *C. tilapiae* (23–34 µ m) reported by Rindoria et al. [102]. The MCO resembles that of *C. tilapiae*. The accessory piece of both species widens distally and terminates in a hook-like structure although, in *C. tilapiae*, the structures curves halfway around the copulatory tube. In *C*. *dibangoi* n. sp., the accessory piece guides the copulatory tube only along the distal portion of the copulatory tube. The copulatory tube of *C*. *dibangoi* n. sp. is also associated with a small heel, which is absent in *C. tilapiae*.

**Name**: *Cichlidogyrus ophioglossa* Moons, Kmentová, Pariselle, Vanhove & Cruz-Laufer n. sp.

**Type-host**: *Chromidotilapia kingsleyae* Boulenger, 1898 (Perciformes: Cichlidae)

**Additional host**: *Chromidotilapia elongata* Lamboj, 1999 (Perciformes: Cichlidae)

**Locality**: small stream, affluent of Moukalaba, Nyanga basin, Gabon; -02.78, 10.77; 19/09/2001; on type host

**Additional locality**: Congo Republic; -4.28, 12.45; on *Chromidotilapia elongata*

**Symbiotype**: RMCA_Vert_2002.006.P.2722-2768 (D)

**Symbioparatype**: RMCA_Vert_1991.068.P.2121-2125 (LA)

**Type-material**: Holotype—RMCA_VERMES_XXXXX, Paratypes—RMCA_VERMES_XXXXX–XXX, HU XXX–XXX, MZH XXXXX–XXX, SAMC XXXXXXX–XXX

**ZooBank registration:** Life Science Identifier (LSID) for *Cichlidogyrus ophioglossa* Moons, Vanhove, Pariselle & Cruz-Laufer n. sp.—urn:lsid:zoobank.org:act:XXXXXXXXXXXX.

**Site in host:** Gills

**Etymology**: The species epithet ‘*ophioglossa*’ is derived from the Greek word *ophis = snake* and *glossa = tongue*, and refers to the morphology of the accessory piece in the male copulatory organ that resembles a forked tongue of a snake.

**Description:** [Based on 76 specimens; Figure 5]. Two pairs of anchors. Ventral anchors with a reduced outer root, inner root more developed. Indentation between roots relatively shallow. Dorsal anchors about the same size as the ventral anchors. Inner root of the dorsal anchor well-developed and outer root reduced. Between the inner and outer root, the anchor shows a small bulge. Ventral transverse bar V-shaped with triangular membranous attachments at distal half of branches. Dorsal transverse bar has thick midsection with two pronounced auricles. Seven pairs of marginal hooks; pairs 1, 3, 4, 5, 6, and 7 with approximately the same length; pair 2 small. Secondary shaft shorter in pair 1 and 4. The MCO consists of a copulatory tube and an accessory piece. Copulatory tube long and slightly curved, narrowing distally, with distal opening; basal bulb broad with heel attached. Accessory piece consisting of two parts, a large distal portion and a proximal connecting piece. The large portion is slightly curved, with a broadened section partly engulfing the copulatory tube. Distal end of the large portion of the accessory piece bifurcating, one end protrudes in a bulbous end, the other end forms a hook with a wing-shaped, serrated structure. The connecting piece is attached at the base of the copulatory tube, bifurcating at the level of the end of the copulatory tube, connecting with the bulbous end of the large portion of the accessory piece. A considerable variation in the MCO morphology is observed in specimens found on the same host individuals (see Figure 6). A string-like structure attaches to the end of the base of the copulatory tube. This attachment point is similar in all individuals. However, the flattening of the specimens during the mounting process results in in different appeareances (see Figure 6 A, B). In some individuals, the string-like structure draws a loop or is curved. In other individuals, this structure is concealed or broken. Hence, the shape of the structure was not always observed. In these cases, the connecting portion is concealed by the large portion of the accessory piece might, which might create the illusion that the large portion is directly connected with the copulatory tube where the string-like structure would attach (see Figure 6C). Furthermore, the wing-shaped structure might appear larger and more open in these individuals (Figure 6C); whereas usually this structure mostly (or partially) overlaps with the large portion of the accessory piece. The sclerotised vagina is tubiform, drawing a U-turn.

**Figure 5:**
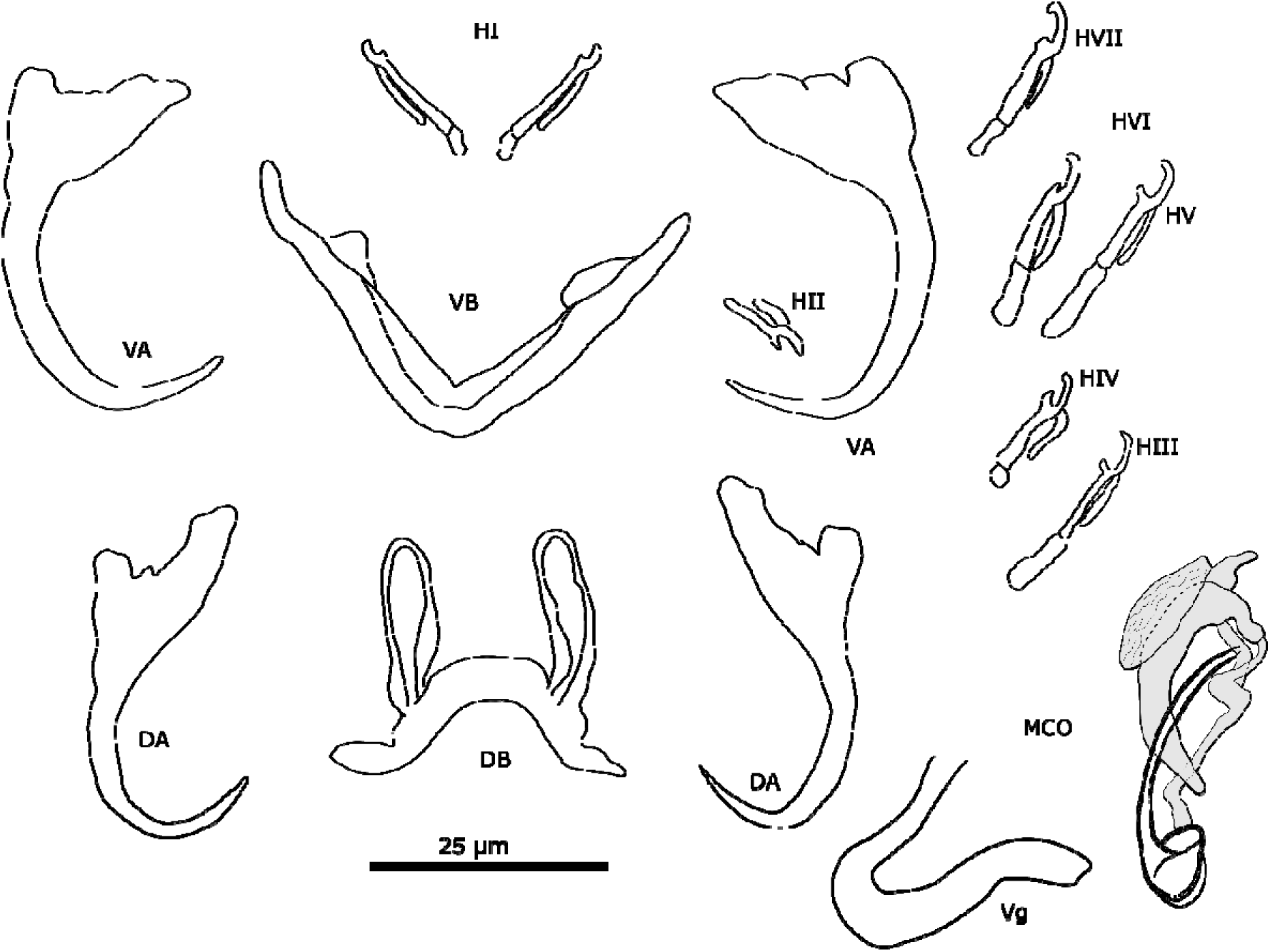
Sclerotised structures of *Cichlidogyrus ophioglossa* n. sp. Abbreviations: HI-HVII, hooks; VA, ventral anchor; VB, ventral transverse bar; DA, dorsal anchor; DB, dorsal transverse bar; MCO, male copulatory organ; Vg, vagina.

**Figure 6:**
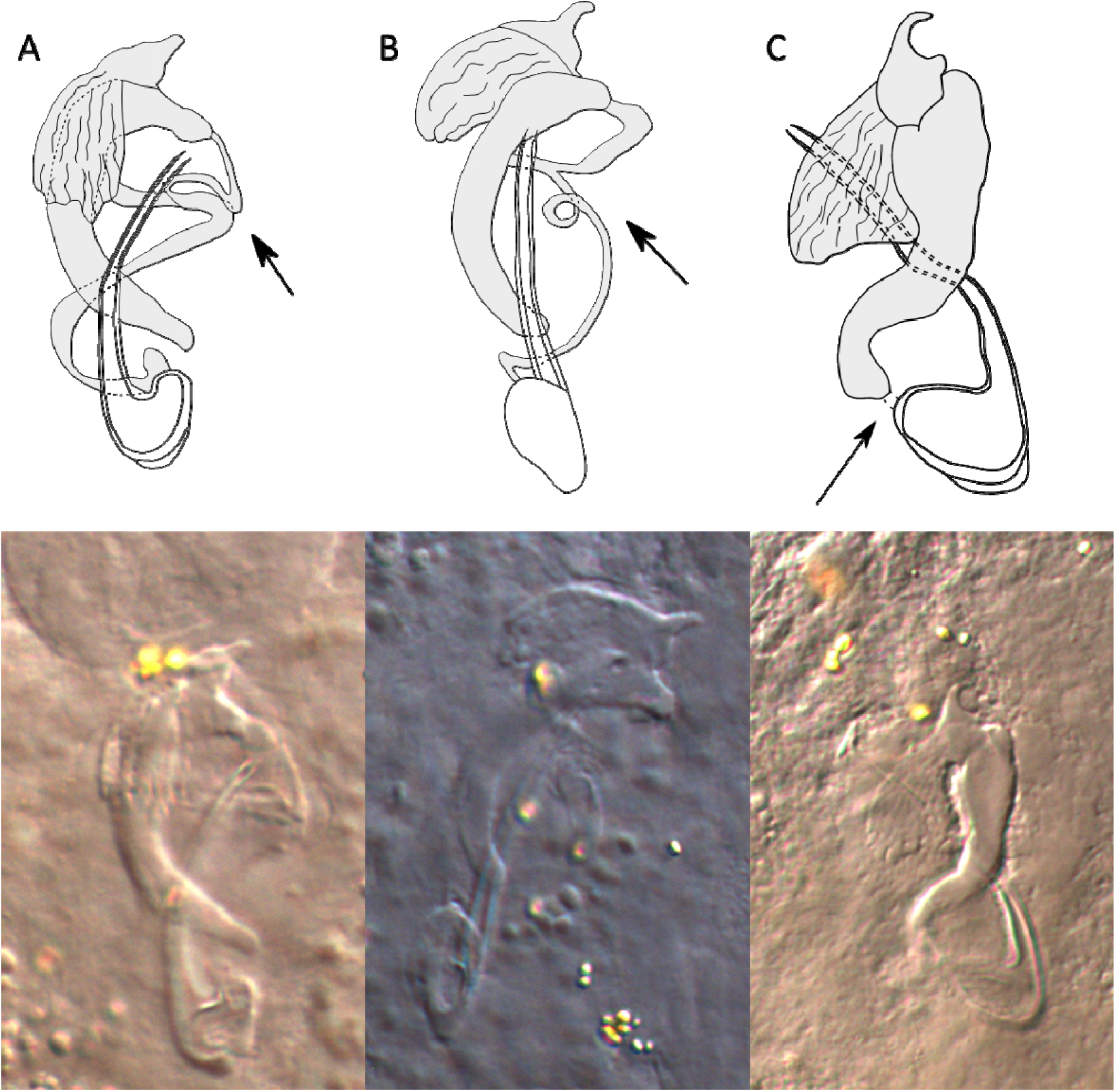
Drawings and microscopic pictures of the male copulatory organs of multiple individuals of *C. ophioglossa* n. sp. Arrows indicating the variation seen in different specimens.

**Remarks**: All specimens show diagnostic features of species of *Cichlidogyrus* (see “Remarks” *C. ataikputu* n. sp.). *Cichlidogyrus ophioglossa* n. sp. resembles *C. acerbus* Dossou, 1982 [14], *C. fontanai* Pariselle & Euzet, 1997 [81], *C. lagoonaris* Paperna, 1969 [74], and *C. nageus* Řehulková, Mendlová & Šimková, 2013 [100]; all infecting *Sarotherodon* species [14,74,81,100]. *Cichlidogyrus acerbus, C. fontanai*, *C. lagoonaris* and *C. nageus* share similarities with *C. ophioglossa* n. sp. in the morphology of the ventral bar. The species have a V-shaped bar with membranous triangles attached at the midsection. The dorsal anchors are also similar in having a well-developed inner root. The ventral anchors of *C. acerbus*, *C. fontanai, C. lagoonaris* and *C. nageus* present distinct roots, where the inner root is more developed than the outer root, while *C. ophioglossa* n. sp. has no distinct roots. The dorsal bar has well-developed auricles in *C. ophioglossa* n. sp. and the other species, yet the midsection of the dorsal bar is thicker in *C. fontanai* (12 µ m) and *C. nageus* (8 µ m) [81, 100], than in *C. ophioglossa* n. sp (5.6 µ m). The copulatory tube in *C. ophioglossa* n. sp. is similar to *C. fontanai*, *C. lagoonaris*, and *C. nageus*, which also have a slightly curved copulatory tube with a broad base. The size of the heel in *C. ophioglossa* n. sp. is as small as observed in *C. fontanai*. The accessory piece of *C. fontanai* is bifurcated at the distal end, which is also seen in *C. ophioglossa* n. sp. Furthermore, a smaller portion of the accessory piece is also observed in *C. nageus*. This part is connected to the broad base of the copulatory tube, as in *C. ophioglossa* n. sp. However, the small portion is string-like in *C. ophioglossa* n. sp., but broader and more finger-like in *C. nageus.* The larger portion of the accessory piece ends in three processes of which one is hook-shaped, also seen in *C. ophioglossa* n. sp. but here a wing-shaped serrated structure is attached to it. No wing-like serrated structure has been reported in any species of *Cichlidogyrus* to date.

**Name**: *Cichlidogyrus gnomon* Moons, Kmentová, Pariselle, Vanhove & Cruz-Laufer n. sp.

**Type-host***: Chromidotilapia elongata* Lamboj, 1999 (Perciformes: Cichlidae)

**Locality**: Mavemba river, tributary of Loukoula on the right bank, 2 km downstream from Mpounga, Republic of the Congo; -4.28, 12.45; 30/07/1991

**Symbiotype**: RMCA_Vert_1991.068.P.2121-2125 (LA)

**Type-material**: Holotype—RMCA_VERMES_XXXXX, Paratypes—RMCA_VERMES_XXXXX–XXX, HU XXX

**ZooBank registration:** Life Science Identifier (LSID) for *Cichlidogyrus gnomon* Moons, Kmentová, Pariselle, Vanhove & Cruz-Laufer n. sp.—urn:lsid:zoobank.org:act:XXXXXXXXXXXX.

**Site in host:** Gills

**Etymology**: The species epithet ‘*gnomon*’ refers to the part of a sundial that casts a shadow. The term is commonly used to refer to an L-shape in geometry. Here, ‘*gnomon*’ refers to the L-shaped accessory piece of the male copulatory organ.

**Description**: [based on 4 specimens, Figure 7] Two pairs of anchors. Ventral anchor with reduced outer root, inner root more developed. Dorsal anchor approximately the same size as the ventral anchor. Outer root of dorsal anchor reduced, slightly larger than the outer root of the ventral anchor. Inner root more developed and larger than the inner root of the ventral anchor. Ventral transverse bar V-shaped with triangular membranous attachments along distal half of branches. Dorsal transverse bar has a thick midsection with auricles. Auricles are drop-shaped. Most likely seven pairs of marginal hooks like all congeners, but pair 2 was not observed due to the poorly preserved specimens. Pairs 1 and 3–7 approximately the same length. Secondary shaft of pairs 1 and 4 shorter. Male copulatory organ consists of a copulatory tube and an accessory piece. Distal opening of the copulatory tube slightly curved. Cupulatory tube narrows towards the distal end and has a broad basal bulb with a small heel. Accessory piece bends in the middle portion and connects to the base of the copulatory tube at two points. Distal end of the accessory piece splits and forms a long and a short projection, each with a bulbous portion. Shorter protrusion connected to a plate. This plate has a hook-like projection, a small bulge at the distal end, and a drop-like projection at the proximal end. No sclerotised vagina observed.

**Figure 7:**
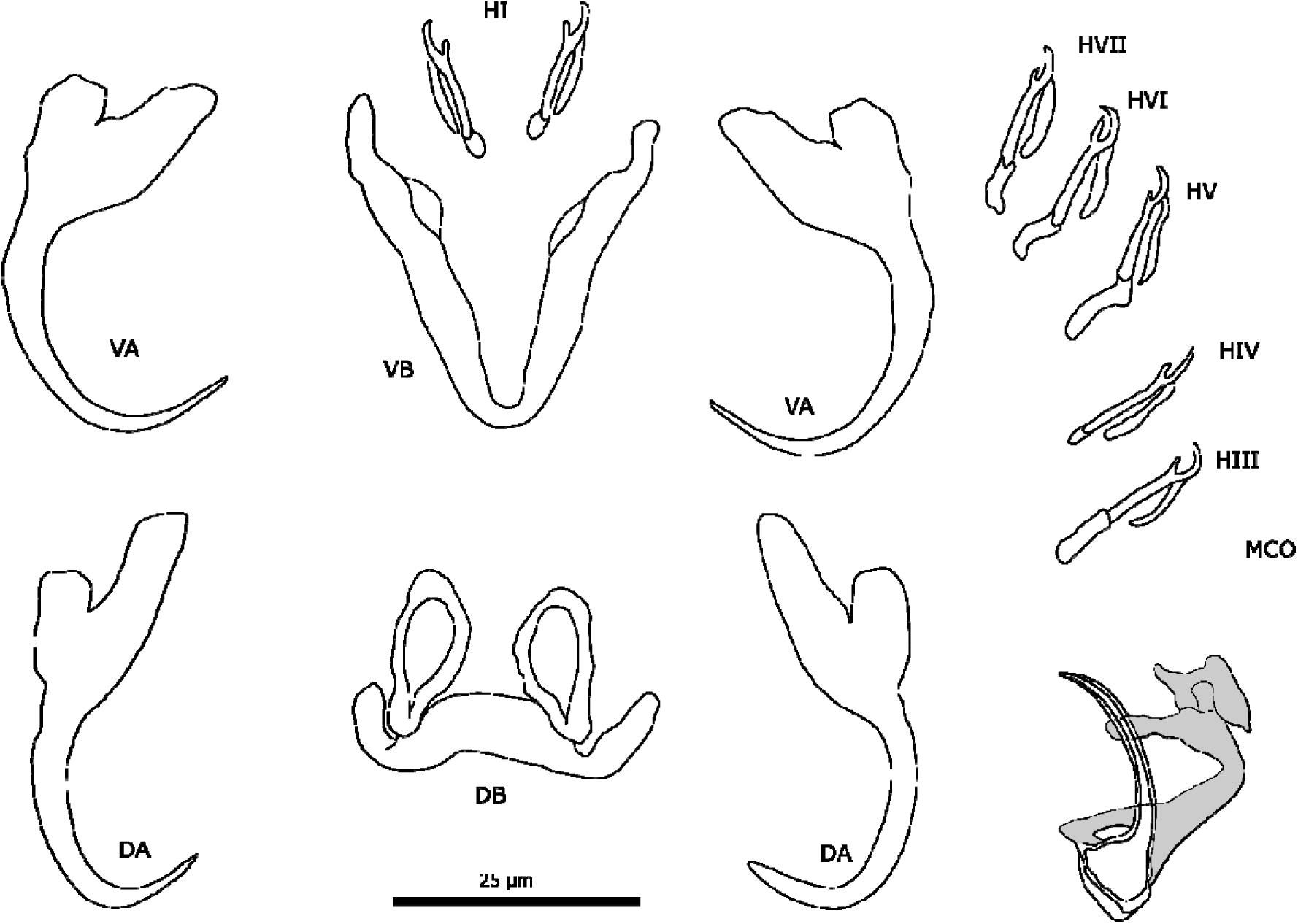
Sclerotised structures of *Cichlidogyrus gnomon* n. sp. Abbreviations: HI-HVII, hooks; VA, ventral anchor; VB, ventral transverse bar; DA, dorsal anchor; DB, dorsal transverse bar; MCO, male copulatory organ.

**Remarks**: All specimens show diagnostic features of species of *Cichlidogyrus* (see “Remarks” *C. ataikputu* n. sp.). *Cichlidogyrus gnomon* n. sp. resembles *C. fontanai* [infecting *Sarotherodon occidentalis* (Daget, 1962) in Bourouma River (Guinea)], in the same way it resembles *C. ophioglossa* n. sp. The protrusions at the distal end of the accessory piece are hook-like in *C. fontanai* but bulbous in *C. gnomon* n. sp. The outer roots of the dorsal anchor are larger in *C*. *gnomon* n. sp. (7.5 µm compared to 2.0 µ m in *C. fontanai*); the inner roots of the ventral anchor (15.0 µ m) are slightly larger than the inner root of the dorsal anchor (11.6 µ m). In *C. fontanai*, the inner roots of dorsal and ventral anchors are approximately the same size. The dorsal transverse bars are similarly shaped, but the dorsal bar is generally larger in *C. fontanai* (DBx = 34 µ m compared to 22 µ m in *C. gnomon* n. sp.). The dorsal bar of *C. muterezii* Pariselle & Vanhove, 2015 [124] resembles *C*. *gnomon* n. sp., but the midsection is thinner in *C. muterezii* (6.4 µ m) than in the former species (13.7 µ m). *Cichlidogyrus gnomon* n. sp. resembles *C. ophioglossa* n. sp in a number of characters. First, the sizes of the ventral and dorsal anchors are similar; morphologically the species differ in the ventral anchors as the incision between the roots is more pronounced in *C. gnomon* n. sp. than in *C. ophioglossa* n. sp. In *C. ophioglossa* n. sp., the accessory piece attaches to the base of the copulatory tube with a small string-like extension whereas, in *C. gnomon* n. sp., the accessory piece attaches to the base of the copulatory tube directly. The accessory piece bifurcates at the distal end for both species but in *C. gnomon* n. sp. this results in two bulbous protuberances. An additional plate-like structure is connected to one of these protuberances, which unlike the wing-shaped structure in *C. ophioglossa* n. sp., is not serrated.

**Name**: *Cichlidogyrus tshuapa* Moons, Kmentová, Pariselle, Vanhove & Cruz-Laufer n. sp.

**Type-host***: Congochromis dimidiatus* (Pellegrin, 1900)

**Locality**: Boende, Tshuapa Province, Boende, Democratic Republic of the Congo; 0.23, 20.83; 29/09/1969

**Symbiotype**: RMCA_Vert_P.174947-174968 (967)

**Type-material**: Holotype—RMCA_VERMES_XXXXX, Paratypes—RMCA_VERMES_XXXXX

**ZooBank registration:** Life Science Identifier (LSID) for *Cichlidogyrus tshuapa* Moons, Kmentová, Pariselle, Vanhove & Cruz-Laufer n. sp.—urn:lsid:zoobank.org:act:XXXXXXXXXXXX.

**Site in host:** Gills

**Etymology**: The species epithet refers to the name of the province Tshuapa, where the species was found. **Description:** [based on 2 specimens, Figure 8] Two pairs of anchors. Ventral anchors with well-developed inner and outer root. Dorsal anchor also with well-developed inner and outer root with the outer root being about half the length of the inner root. Both dorsal and ventral anchors have deep indentations. Ventral transverse bar V-shaped with membranous attachments over most of the length of the branches. Dorsal transverse bar with thickened midsection and auricles. Seven pairs of marginal hooks. The first pair is large (‘standardized length’ larger than 1.7 following Pariselle & Euzet [79]) with long and broad secondary shafts. Pairs 3–7 approximately the same length. Pair 2 is the smallest. The male copulatory organ consists of a copulatory tube, an accessory piece, and a heel. The copulatory tube has a broad middle section and narrows at the distal end. The S-shaped accessory piece is attached to the heel, which is small. The distal portion is positioned parallel to the copulatory tube. The sclerotised vagina is pear-shaped with a tubiform sinuous extension.

**Figure 8:**
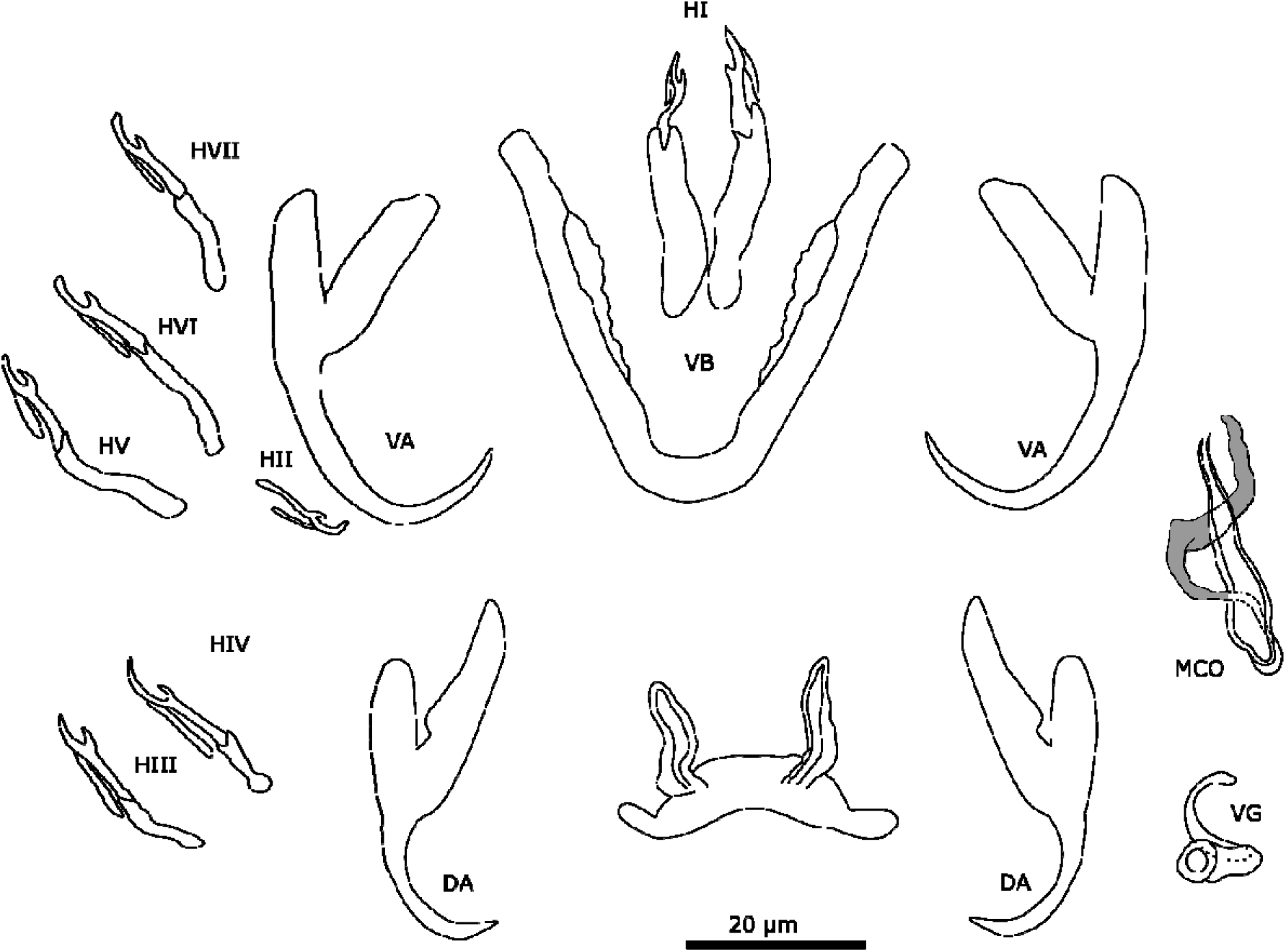
Sclerotised structures of *Cichlidogyrus tshuapa* n. sp. Abbreviations: HI-HVII, hooks; VA, ventral anchor; VB, ventral transverse bar; DA, dorsal anchor; DB, dorsal transverse bar; MCO, male copulatory organ.

**Remarks**: All specimens show diagnostic features of species of *Cichlidogyrus* (see “Remarks” *C. ataikputu* n. sp.). The new species shows strong resemblances with *Cichlidogyrus papernastrema* Price, Peebles & Bamford, 1969 [94] infecting *Tilapia sparrmanii* Smith, 1840 [94], *Oreochromis mweruensis* Trewavas, 1983, and *Coptodon rendalli* (Boulenger, 1897). The copulatory tube of *Cichlidogyrus tshuapa* n. sp. has a slightly broadened middle section, whereas, in *C. papernastrema*, this section forms a more apparent rounded bulb. In the redescription of *C. papernastrema* by Jorissen et al. [42], a heel is present and the accessory piece is described as S-shaped. The curvature of the accessory piece at the distal end is almost the same as in *C. tshuapa* n. sp. However, the curvature of the accessory piece in *C. tshuapa* n. sp. is sharper than in the original description [94] and the redescription [42] of *C. papernastrema*. Furthermore, the haptor differs substantially. The outer root of the dorsal anchors is larger in *C. tshuapa* n. sp. (on average 13.3 µ m compared to 7 µ m in *C. papernastrema* [94]), as is the inner root (25.4 µ m compared to 17 µ m). The ventral bar of *C. papernastrema* also lacks membranous extensions unlike in *C. tshuapa*. Furthermore, *C. tshuapa* n. sp. presents a sclerotised vagina unlike *C. papernastrema*.

**Name**: *Cichlidogyrus thysochromi* Moons, Kmentová, Pariselle, Vanhove & Cruz-Laufer n. sp.

**Host**: *Thysochromis ansorgii* (Boulenger, 1901)

**Locality**: Aboisso, Côte d’Ivoire; 4.47, -3.2; November 1958

**Symbiotype**: RMCA_Vert_1973.005.P.4470-4476 (447)

**Type-material**: Holotype—RMCA_VERMES_XXXXX

**ZooBank registration:** Life Science Identifier (LSID) for *Cichlidogyrus thysochromi* Moons, Vanhove, Pariselle & Cruz-Laufer n. sp.—urn:lsid:zoobank.org:act:XXXXXXXXXXXX.

**Site in host:** Gills

Etymology: The species epithet refers to the host genus *Thysochromis*, representatives of which the species was found on.

**Description**: [based on 1 specimen, Figure 9] Two pairs of anchors. Ventral anchor with a well-developed inner root and smaller outer root. Dorsal anchors present but distorted on slide. Ventral transverse bar V-shaped. Membranous attachments attached over the length of the bar. Dorsal transverse bar with thick midsection and auricles. The auricles are small and slender. Most likely seven pairs of marginal hooks like all congeners, but pair 3 was not observed due to the poorly preserved specimen. The first pair is larger than the other hooks. The rest of the hooks are about the same size. The male copulatory organ consists of a copulatory tube, accessory piece, and a heel. The base of the copulatory tube is broad. The copulatory tube makes a 90° turn near the base and narrows distally. The heel curves around the base of the copulatory tube and is broad. The accessory piece is shaped drop-like and surrounds the copulatory tube distally. The basal portion of the accessory piece was not observed. No sclerotised vagina was observed.

**Figure 9:**
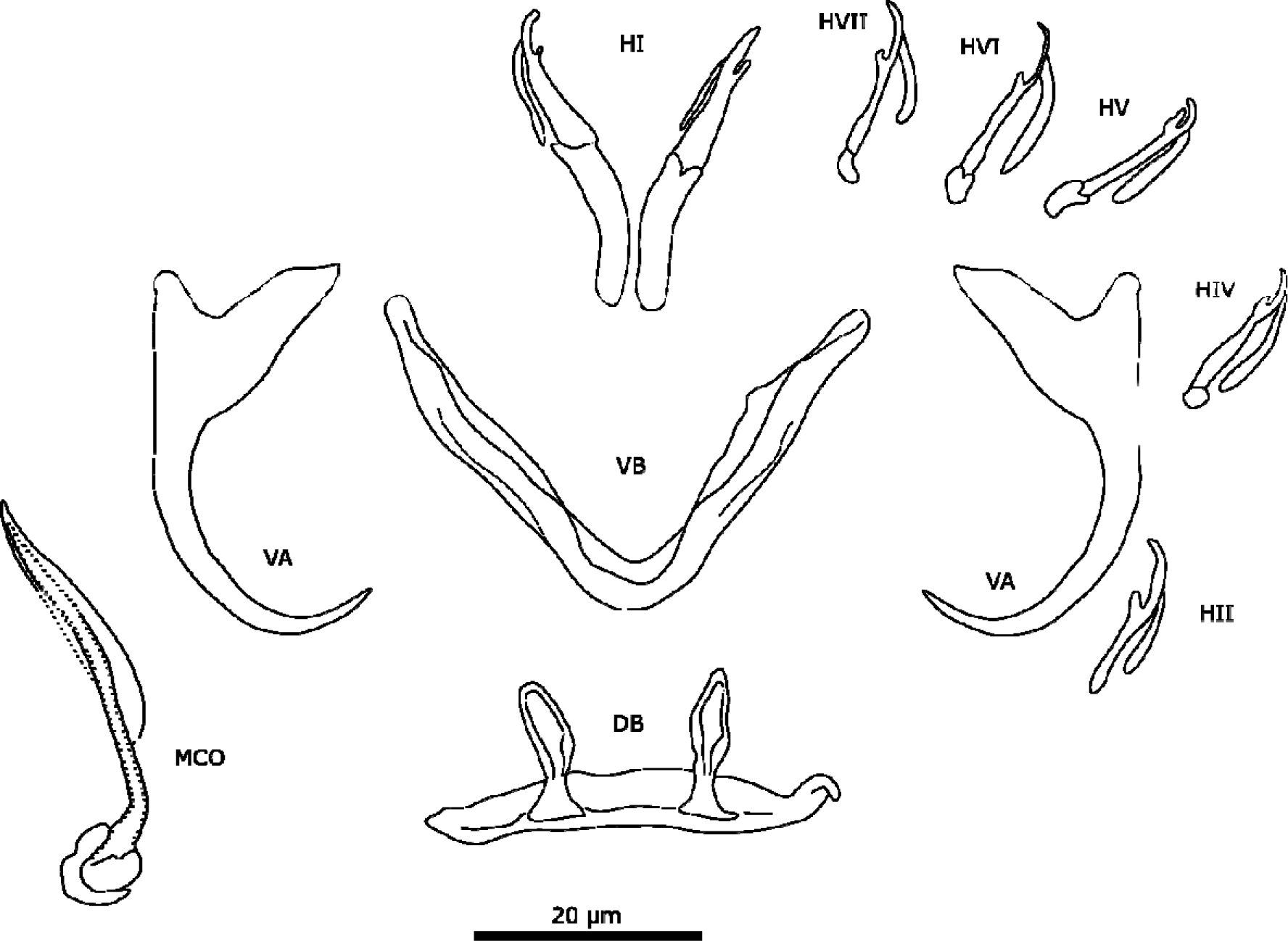
Sclerotised structures of *Cichlidogyrus thysochromi* n. sp. Abbreviations: HI-HVII, hooks; VA, ventral anchor; VB, ventral transverse bar; DA, dorsal anchor; DB, dorsal transverse bar; MCO, male copulatory organ.

**Remarks**: The only observed specimen shows diagnostic features of species of *Cichlidogyrus* (see “Remarks” *C. ataikputu* n. sp.). Although multiple fish were examined for parasites, only one specimen of this species was found. While describing a new species based on single individuals (singletons) is unusual for monogeneans, previous studies have done so with partial specimens or a few specimens when the morphology was discernible and distinct from other species [43] and the practice is widespread among other taxa, e.g. arthropods [58]. Therefore, we opted to describe the present specimen as a new species. *Cichlidogyrus thysochromi* n. sp. does show some similarities, e.g. in the haptor, with *Cichlidogyrus polyenso* Jorissen, Pariselle & Vanhove, 2018 [43] and *Cichlidogyrus calycinus* Kusters, Jorissen, Pariselle & Vanhove, 2018 [43] both infecting *Hemichromis elongatus* (Guichenot, 1861), *Cichlidogyrus teugelsi* Pariselle & Euzet, 2004 [82] infecting *Hemichromis fasciatus* Peters, 1858, and *Cichlidogyrus reversati* Pariselle & Euzet, 2003 [77] infecting *Pelmatolapia cabrae* (Boulenger, 1899). The ventral anchors of the different species have a well-developed inner roots and small outer roots. The morphology of the ventral bar of *C. polyenso* resembles the one of *C*. *thysochromi* n. sp. the most by having a membranous attachment associated with the ventral bar. The auricles of the dorsal transverse bar are of similar size as the auricles of species mentioned above. The size of the auricles falls within the range of *C. calycinus*, *C. teugelsi*, and *C. polyenso*. Pair 1 of the marginal hooks is also similar in the morphology and size in having a broad and long secondary shaft. The size of the marginal hook pair 1 of *C*. *thysochromi* n. sp. falls within the range of *C. teugelsi* and *C. reversati*. Conversely, the MCO of *C. thysochromi* n. sp. has no resemblances to the MCO of the above mentioned species. In all species the copulatory tube is G-shaped (*C. calycinus* and *C. teugelsi*) or spiralled (*C. polyenso*), while in *C. thysochromi* n. sp. the copulatory tube is slightly curved.

**Name**: Cichlidogyrus sp. ‘Pelvicachromis roloffi’

**Host**: Pelvicachromis roloffi Paperna, 1968

**Locality**: Kahmranka, near Rokupr 10-15 km, route Rokupr-Kambia, Sierra Leone; 9.07, -12.93; 5/4/1969

**Host voucher**: RMCA_Vert_P.174947-174968 (A)

**Parasite material**: RMCA_VERMES_XXXXX

**Description**: [based on 1 specimen, Figure 10] The male copulatory organ consists of a copulatory tube and an accessory piece. The copulatory tube is straight with no heel attached to the base. The distal end of the copulatory tube is not observed as the accessory piece is folded over its distal portion. The accessory piece shows a minor split at the distal end and is attached to the base of the copulatory tube.

**Figure 10:**
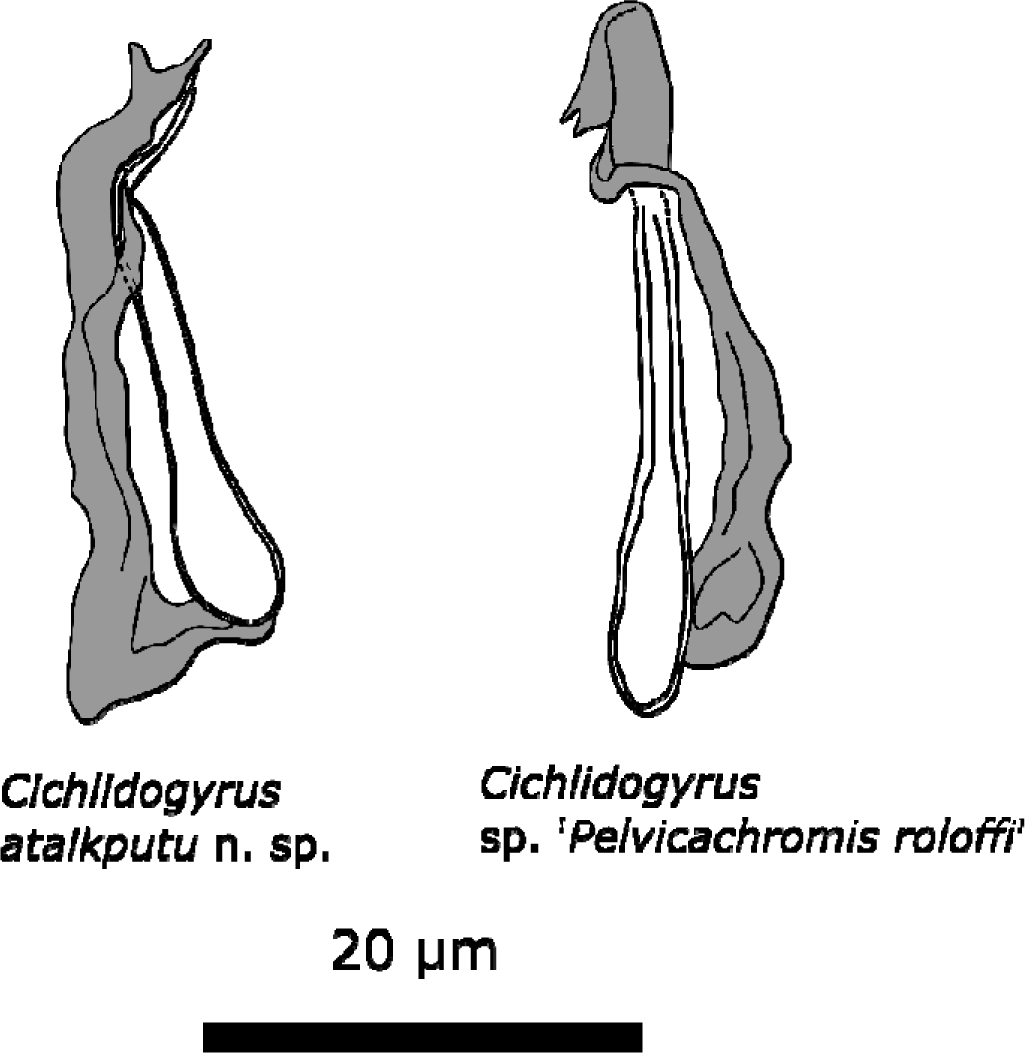
Comparison of the MCO of Cichlidogyrus ataikputu and Cichlidogyrus sp. ‘Pelvicachromis roloffi’.

**Remarks**: Only one specimen was found on the gills of *Pelvicachromis roloffi* but the haptor was lost during the sampling process and the MCO appeared slightly distorted resulting from the fixation on the slide. Based on this information, the specimen could not be assigned to a species or delimited from other species. The MCO resembles the MCO of *Cichlidogyrus ataikputu* n. sp. The fold over the copulatory tube is seen in *C. ataikputu* n. sp. and the split at the distal end is also present in *C. ataikputu* n. sp. The average size of the copulatory tube of *C. ataikputu* is 25.0 µ m (± 4.1 µ m) and the average length of the accessory piece is 27.2 µ m (± 5.8 µ m); measurements of *C*. sp. ‘*Pelvicachromis roloffi*’ are respectively 28.7 and 29.3 µ m. These measurements fall within the range of *C. ataikputu* n. sp. However, the different attachment point of the accessory piece to the basal bulb of the copulatory tube strongly suggests that *C*. sp. ‘*Pelvicachromis roloffi*’ is a new species. However, there is currently too little information for a formal description.

Subclass Polyonchoinea Bychowsky, 1937

Order Dactylogyridea Bychowsky, 1937

Dactylogyridae Bychowsky, 1933

*Onchobdella* Paperna, 1968

**Emended diagnosis** (based on Paperna [72] and Pariselle & Euzet [83]): Body shape elongated or stout. Prohaptoral anterior lobes are poorly demarcated and head organs are present. Two pairs of eyespots that are arranged in pairs of two in front of the pharynx or in one transverse row in front of the pharynx. Three pairs of cephalic glands. Intestinal crura are united posteriorly. The single testis is in the posterior position within the intestinal loop, while the single ovary is located anteriorly to the testis. The MCO consists of a tubiform cirrus and an accessory piece, consisting of two elongated bracket shaped portions. The portions are attached to each other along one or two edges and the copulatory tube often is protruded between. Seminal vesicles and 2–3 prostate glands follow the distal part of the male genital system. Sclerotised vagina present, opening lateral. Anchors two pairs unequal in size and shape. One dorsal, large pair and one ventral small pair more hook shaped. Three transversal bars, one sturdy dorsal bar that is either slightly curved or horse-shoe shaped, and a ventral bar split in two, frequently curved bars. Each anchor is accompanied by membranous filaments, attached to the shaft. Seven pairs of marginal hooks all approximately the same length with thin needle-shaped shaft with delicate posterior projecting process.

**Remarks**: Species of *Onchobdella* are reported from *Chromidotilapia guntheri* (Sauvage, 1882), *Pelmatochromis buettikofferi* (Steindachner, 1894), and species of *Hemichromis* Peters, 1857; and are mainly found in West and Central Africa. The genus was created in 1968 to include five new species (*O. aframae* Paperna, 1968, *O. krachii* Paperna, 1968 *O. pterigyalis* Paperna, 1968, *O. spirocirra* Paperna, 1968, and *O. voltensis* Paperna, 1968 [72]). Paperna considered the presence of two pairs of anchors of unequal size, an accessory piece consisting of two elongated bracket shaped portions, and having three transversal bars as the main characteristics in the first diagnosis [72]. Since Paperna’s diagnosis, six new species (*O. bopeleti* Bilong Bilong & Euzet, 1995 [3], *O. melissa* Pariselle & Euzet, 1995 [83], *O. sylverai* Pariselle & Euzet, 1995 [83], and *O. ximenae* Jorissen, Pariselle & Vanhove in Jorissen et al. [43]) were described, including the two new species described here. As several characteristics deviate from the original diagnosis, we provide an emended diagnosis here. In the original diagnosis, the dorsal bar was described as a frequently curved bar. The first species of *Onchobdella* that were described had either horseshoe-shaped or slightly curved dorsal bars [72]. The dorsal bar is horseshoe-shaped in *O. voltensis*, *O. spirocirra*, *O*. *pterigyalis* and *O. bopeleti* unlike the slightly curved dorsal bar of *O. aframae*, *O. krachii*, *O. melissa* and *O. silverai*. The ventral bar is split in two. Two pairs of eyes are observed in *O. macrohamulus* n. sp., *O. yemojae* n. sp., *O. silverai* and *O. melissa* [83]. Lastly, 14 marginal hooks are counted in the species described after Paperna [72], while the original diagnosis mentions only 4 to 6 pairs of marginal hooks. Pariselle and Euzet [83] already remarked on this difference. Notably, the hooks are difficult to count correctly as the large dorsal anchors often conceal their presence.

**Name**: *Onchobdella macrohamulus* Moons, Kmentová, Pariselle, Vanhove & Cruz-Laufer n. sp.

**Type-host***: Thysochromis ansorgii* (Boulenger, 1901) (Perciformes: Cichlidae)

**Locality**: Oshika, 10 km NW of Ahoada, Nigeria; 5.12, 6.63; March 1984

**Additional locality**: Attingué, Agnébi basin, Côte d’Ivoire; 5.47, -4.183 and Dahomey, Whedda, Ouémé river, Benin, 6.75, 2.457 on *Thysochromis ansorgii*

**Symbiotype:** RMCA_Vert_1984.022.P.0012-0014 (A)

**Symbioparatype:** RMCA_Vert_1973.005.P.4470-4476 (447); RMCA_Vert_1973.005.P.4478-4503 (502)

**Type material**:

**Type-material**: Holotype—RMCA_VERMES_XXXXX, Paratypes—RMCA_VERMES_XXXXX–XXX, HU XXX, SAMC XXXXXXX

**ZooBank registration:** Life Science Identifier (LSID) for *Onchobdella macrohamulus* Moons, Vanhove, Pariselle & Cruz-Laufer n. sp.—urn:lsid:zoobank.org:act:XXXXXXXXXXXX.

**Site in host:** Gills

**Etymology**: The species epithet is a combination of the Greek word ‘macro’ (= long) and the Latin word ‘hamulus’ (= hook-shaped carpal bone). The combination is used to describe the exceptionally large dorsal anchors for a species of *Onchobdella*.

**Description:** [based on 5 specimens, Figure 11] Dorsal anchors are very large. Outer root of the large anchor is reduced while inner root is well-developed. The outer roots of the small (ventral) anchors are more developed, but still smaller the inner roots. Dorsal bar stout and straight, slightly curved at both ends. Ventral bars slightly curved. Seven pairs of hooks all approximately the same length and thin, needle-like. Male copulatory organ consists of copulatory tube and accessory piece. Copulatory tube strongly curved with a broad, bean-shaped base. At the level of the accessory piece, the copulatory tube is almost straight. The base of the accessory piece connects to the base of the copulatory tube. The accessory piece consists of two portions that connect to each other at the base. The tips of the two portions are slightly curved at the distal end and do not connect with each other. No sclerotised vagina observed.

**Figure 11:**
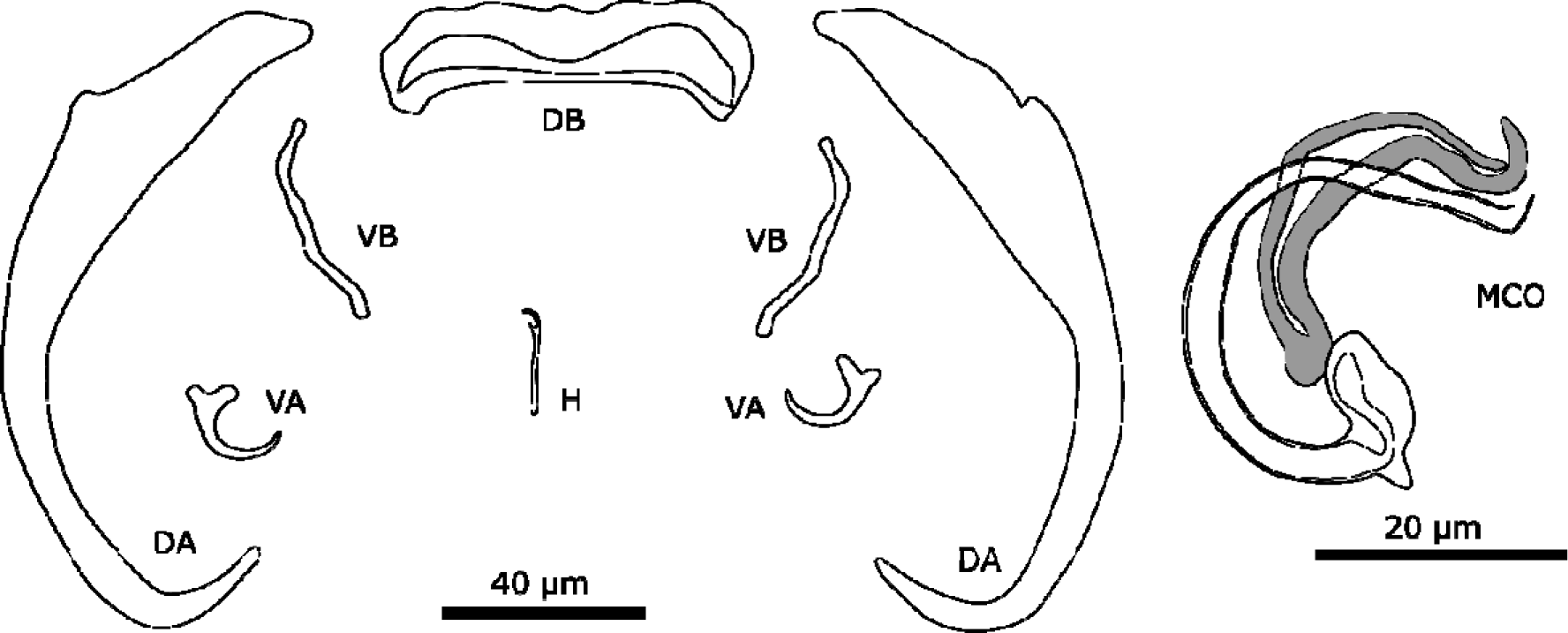
Sclerotised structures of *Onchobdella macrohamulus* n. sp. Abbreviations: H, marginal hook; VA, ventral anchor; VB, ventral transverse bar; DA, dorsal anchor; DB, dorsal transverse bar; MCO, male copulatory organ.

**Remarks:** The specimens show typical features of members of *Onchobdella*: (i) two pairs of anchors unequal in size, one larger and one smaller pair; (ii) two ventral bars, club shaped; (iii) one dorsal bar, curved solid; (iv) hooks seven pairs, needle-shaped [81]. *Onchobdella macrohamulus* n. sp. shows similarities with *Onchobdella krachii* Paperna, 1968 infecting *Chromidotilpia guntheri* (Sauvage, 1882) and *Chromidotilapia linkei* Staeck, 1980. The species has exceptionally large dorsal hooks (DAa on average: 120.0 µ m), three times the length of those of *O. yemojae* n. sp. (Table 5), which is found on the same host species (see description below). Other components of the haptor are in the size range that is known from previously described species of *Onchobdella*. The dorsal bar of *O. macrohamulus* n. sp. different from those of *O. krachii*, whose dorsal bars are broad and straight with curved edges. The morphology of the dorsal anchor is different from *O. krachii*. The outer root is almost not developed in *O. macrohamulus* n. sp. unlike in *O. krachii*. Ventral bars of *O. macrohamulus* n. sp. are more curved than the ones of *O. krachii*. The two portions of the accessory piece are, however, not connected at the distal end in *O. macrohamulus* n. sp. unlike in the other two species. In *O. krachii*, the base of the copulatory tube is round.

**Name**: *Onchobdella yemojae* Moons, Kmentová, Pariselle, Vanhove & Cruz-Laufer n. sp.

**Type-host***: Thysochromis ansorgii* (Boulenger, 1901) (Perciformes: Cichlidae)

**Locality**: Dahomey, Whedda, Ouémé river, Benin; 6.75, 2.47; 14/10/1966

**Additional locality**: Attingué, basin Agnébi, Côte d’Ivoire; 5.47, -4.183 and Oshika, 10km NW of Ahoada, Nigeria; 5.117, 6.633 on *Thysochromis ansorgii*

**Symbiotype**: RMCA_Vert_1973.005.P.4478-4503 (447)

**Symbioparatype**: RMCA_Vert_1973.005.P.4470-4476 (500, 502, 503); RMCA_Vert_1984.022.P.0012-0014 (A)

**Type-material**: Holotype—RMCA_VERMES_XXXXX, Paratypes—RMCA_VERMES_XXXXX–XXX, HU XXX–XXX, MZH XXXXX–XXX, SAMC XXXXXXX–XXX

**ZooBank registration:** Life Science Identifier (LSID) for *Onchobdella yemojae* Moons, Vanhove, Pariselle & Cruz-Laufer n. sp.—urn:lsid:zoobank.org:act:XXXXXXXXXXXX. **Site in host**: Gills

**Etymology**: The species epithet is based on the name of the water spirit Yemoja from the Yoruba religion. This religion originates in the countries where *O. yemojae* n. sp. is found but has since then spread to other parts of the world as a result of the Yoruba diaspora.

**Description**: [based on 29 specimens, Figure 12] Two pairs of anchors. The dorsal and ventral anchors have a well-developed inner root and a reduced outer root. The ventral bars are thin and slightly curved, the middle portion is straight with both ends being curved. The dorsal bar curves in the middle and the ends are straight. Seven pairs of thin marginal hooks of similar size. The MCO consists of a copulatory tube and an accessory piece. The copulatory tube has a bean-shaped basal bulb and narrows towards the distal end. The accessory piece consists of two plates that connect at the proximal end, where it also attaches with the base of the copulatory tube.

**Figure 12:**
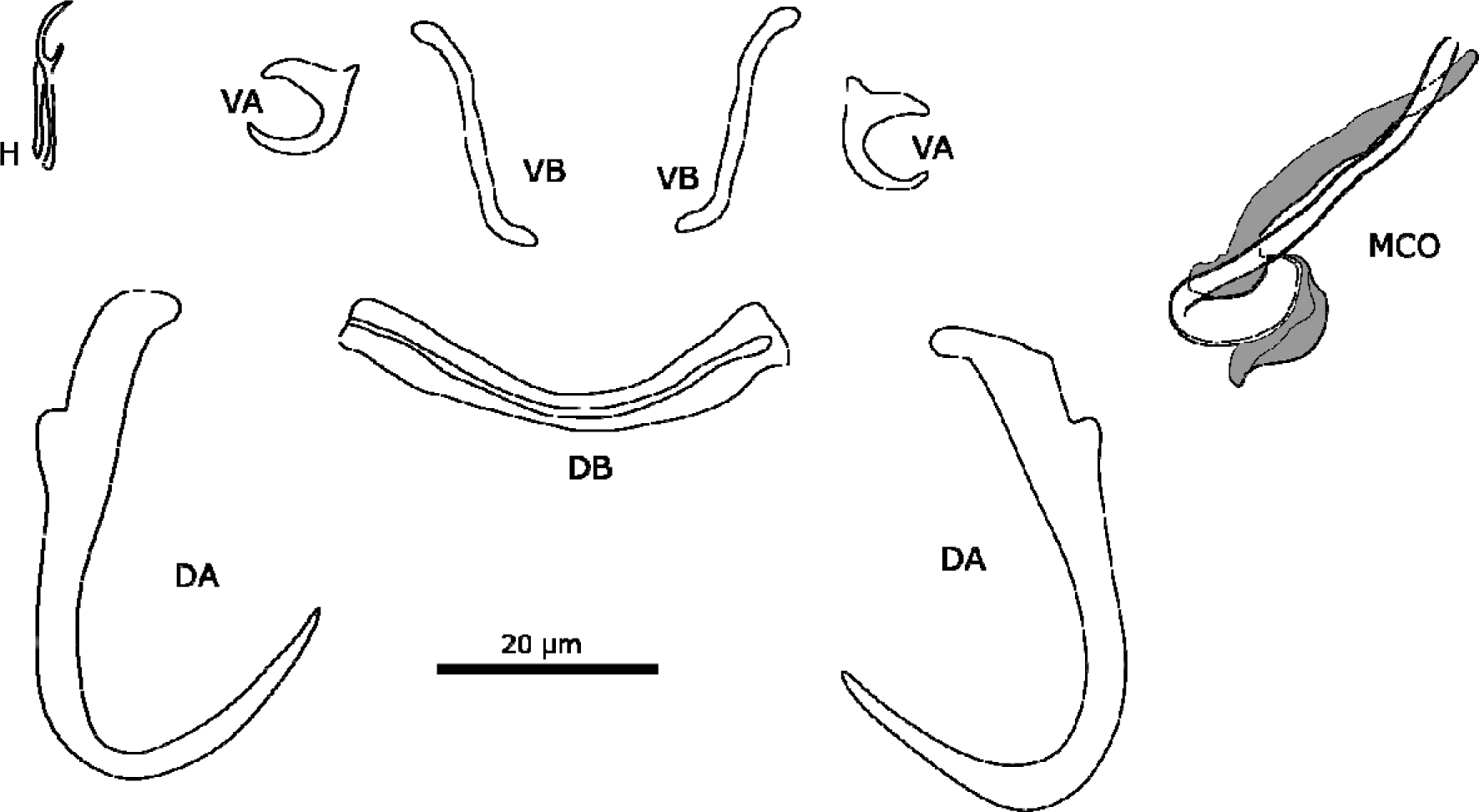
Sclerotised structures of *Onchobdella yemojae* n. sp. Abbreviations: H, marginal hook; VA, ventral anchor; VB, ventral transverse bar; DA, dorsal anchor; DB, dorsal transverse bar; MCO, male copulatory organ.

**Remarks**: All specimens show diagnostic features of species of *Onchobdella* (see remarks on *O. macrohamulus* n. sp.). *Onchobdella yemojae* n. sp. resembles *Onchobdella melissa* Pariselle & Euzet, 1995 infecting *Pelmatochromis buettikoferi* (Steindachner, 1894). The morphology of the haptor of both species are similar. For instance, the dorsal bar is curved slightly with straight ends. However, *O*. *yemojae* n. sp. shows a ridge along the dorsal bar similar to *O. krachii* whereas *O. melissa* lacks this ridge. In *O. krachii*, the bar has bent instead of straight ends. The morphology of the anchors is also similar as both *O. melissa* and *O. yemojae* n. sp. have reduced outer roots and well-developed inner roots. Furthermore, the ventral bars present rounded, slightly bent ends in both *O. krachii*and *O. yemojae*. The accessory piece of *O*. *yemojae* n. sp. is connected to the base of the copulatory tube. In *O. melissa*, the accessory piece is not connected to the base of the copulatory tube and the plates of the accessory piece are of different lengths. The copulatory tube of *O. melissa* is G-shaped, while the copulatory tube of *O. yemojae* n. sp. is J-shaped. The copulatory tube in *O. melissa* is also longer (69 µ m ± 2.9 µm [83]) than in *O. yemojae* n. sp. (36.6 µ m ± 4.7 µ m). The typical structure, two portions connected at the base (accessory piece), is difficult to see in the individuals of *O*. *yemojae* n. sp. as only one of the plates is visible in most cases.

**Name**: Onchobdella krachii Paperna, 1968

**Type locality**: Kpandu and Kete Krachi, Volta Lake, Ghana

**Localities**: New Calabar river, Akpor, Nigeria; 4.87, 6.9; 01/12/1990 on *Chromidotilapia guntheri* (Sauvage, 1882); road Yabassi–Yingui, Cameroon; 07/11/1990 on *Chromidotilapia linkei*

**Host vouchers:** RMCA_Vert_XXXXXX RMCA_Vert_1991.010.P.0542-0582 (576, 577, 578, 580, 581, 582); RMCA_Vert_1992.144.P.0250-0261 (A, B, C)

**Parasite material:** RMCA_VERMES_XXXXX–XXX, HU XXX–XXX, MZH XXXXX–XXX, SAMC XXXXXXX–XXX

**Site on host:** Gills

**Redescription**: [based on 82 specimens, Figure 13] Two pairs of anchors. The dorsal anchors have well-developed inner roots and reduced outer roots. The ventral anchors have well-developed inner roots and small outer roots. The dorsal bar is slightly curved at both ends of the bar. The ventral bars are also slightly curved with at one end a small indentation. Seven pairs of marginal hooks present and all approximately of the same size, thin and needle-shaped. The male copulatory organ consists of a copulatory tube and an accessory piece. The copulatory tube draws a spiral in the shape of the letters G and has a oval-shaped base. The accessory piece is attached to the base of the copulatory tube and consists of two plates that connect at the proximal end. At the distal end, a structure is connected to one of the plates running in parallel with the copulatory tube.

**Figure 13:**
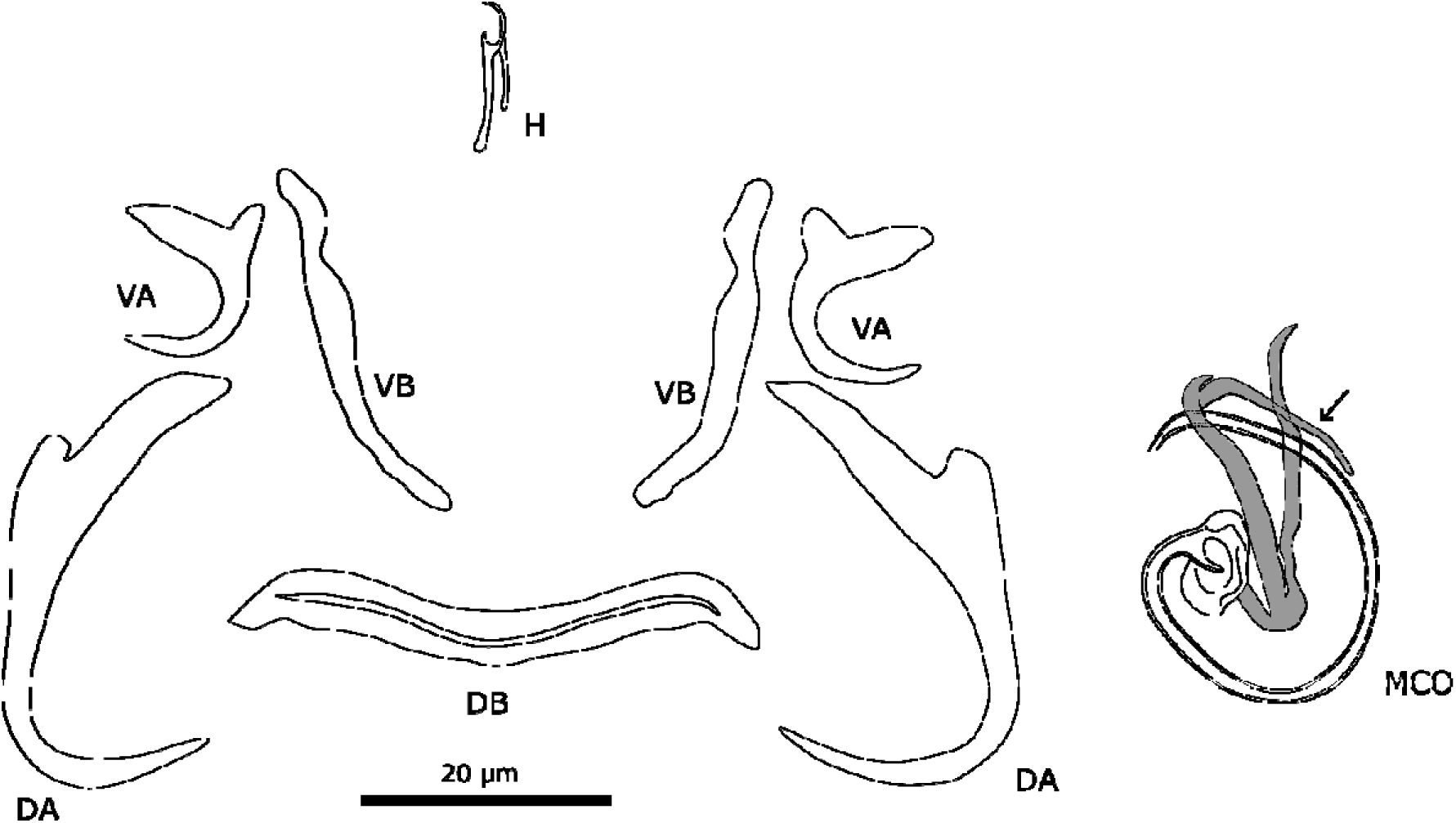
Sclerotised structures of *Onchobdella krachii*. Abbreviations:H, marginal hook; VA, ventral anchor; VB, ventral transverse bar; DA, dorsal anchor; DB, dorsal transverse bar; MCO, male copulatory organ. Arrow indicates additional structure of the MCO that was missing from previous characterizations.

**Remarks**: All specimens show diagnostic features of species of *Onchobdella* (see remarks on *O. macrohamulus* n. sp.). The original description by Paperna [72] was based on two specimens. Here we studied 82 specimens and found additional characteristics. The small indentations in the ventral bars were not reported before. Furthermore, the accessory piece of the MCO differs in small characteristics from the original description. The two plates do not connect with the copulatory tube unlike originally reported. Also, an additional structure has been discovered that is connected to one of the plates (Figure 13).

### Morphometrics

We produced three plots for the principal component analysis (PCA) to test whether the qualitative distinctions between the species infecting chromidotilapiine cichlids translate into morphometric differences between their attachment and reproductive organs. We conducted one PCA including all species of *Onchobdella*sampled here, one with species similar to *Cichlidogyrus tilapiae* including *C. tilapiae* ex *Chromidotilapia* spp., *C. dibangoi* n. sp., and *C. ataikputu* n. sp., and one with all species of *Cichlidogyrus* sampled in the present study (Figure 14).

**Figure 14:**
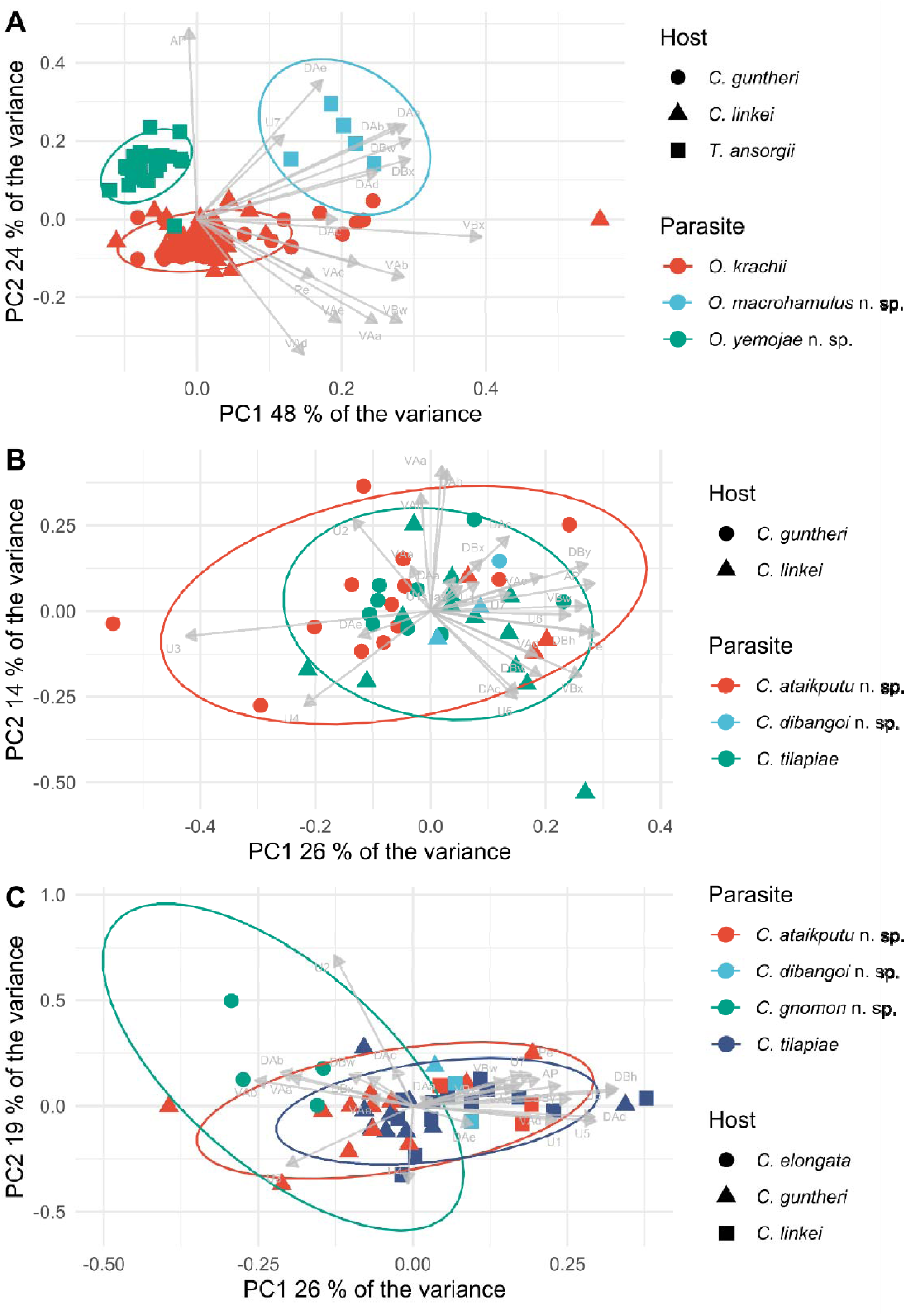
Principal component analyses of monogenean flatworms infecting chromidotilapiine cichlids. A, species of *Onchobdella* showing three distinct clusters that are mostly congruent with the species identities assigned in this study. B, Several species of *Cichlidogyrus* strongly resemble *C. tilapiae*, but at least two of them form distinct species. C, Species of *Cichlidogyrus* infecting chromidotilapiine cichlids form distinct clusters in the PCA, albeit with some overlap.

Specimens belonging to *Onchobdella* (first two principal components together explaining overall 72% of the variation) confirmed the differentiation of the three chromidotilapiines-infecting species. We found two well-separated clusters, with *Onchobdella macrohamulus* n. sp. differing substantially from the other species in the size of the dorsal anchor and the dorsal bar. The clusters produced by specimens of *Onchobdella krachii*and *O. yemojae* n. sp. overlap slightly but with a visible difference in the measurements of the dorsal anchor (Figure 14A), one of the characters we highlighted in the description of *O. macrohamulus* n. sp.

*Cichlidogyrus tilapiae* on one side and the *C. ataikputu* n. sp. and *C. dibangoi* n. sp. on the other side present no apparent clustering (first two PCs explaining 40% of the variation) (Fig. 14B). The pattern indicates that *C. tilapiae* is indistinguishable from the other two species based purely on morphometric characters, highlighting the importance of the qualitative characters used in this study. Specimens of *C. tilapiae* found in this study (Figure 15) were not morphologically distinguishable from specimens from previous studies (as redescribed in [117]) although the measurements differed considerably. Despite the difference in size, we still treat these specimens as belonging to *C. tilapiae* because of a lack of qualitative morphological differences. Finally, when comparing species of *Cichlidogyrus* infecting species of *Chromidotilapia* (first two PCs explaining 47% of the variation), we did not detect any apparent clusters (Fig. 14C).

**Figure 15:**
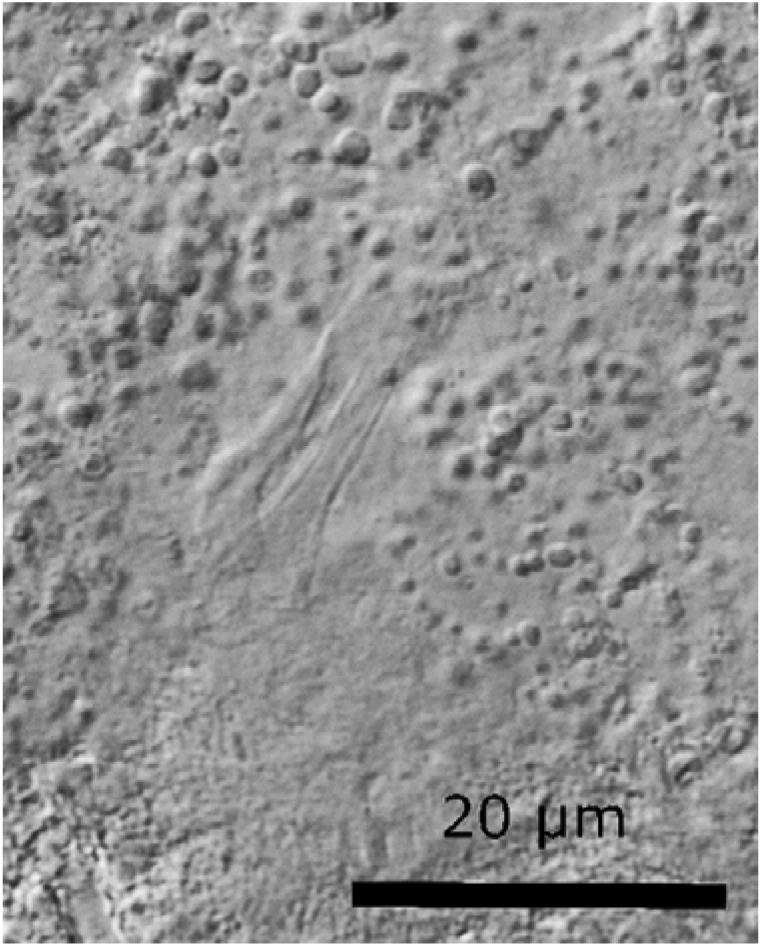
Male copulatory organ (MCO) of a specimen of *Cichlidogyrus tilapiae* infecting *Chromidotilapia guntheri* from Lake Barombi-Kotto.

### Phylogenetic position

According to the parsimony analysis, the species of *Cichlidogyrus* infecting chromidotilapiine cichlids form a well-supported monophyletic group (GC = 95) including specimens identified as *Cichlidogyrus tilapiae* (Figure 16). This group appears firmly nested inside a well-supported (GC = 31) clade of Western African species known to infect mostly hemichromine cichlids, see ‘*Hemi*’ group *sensu* [12].

**Figure 16:**
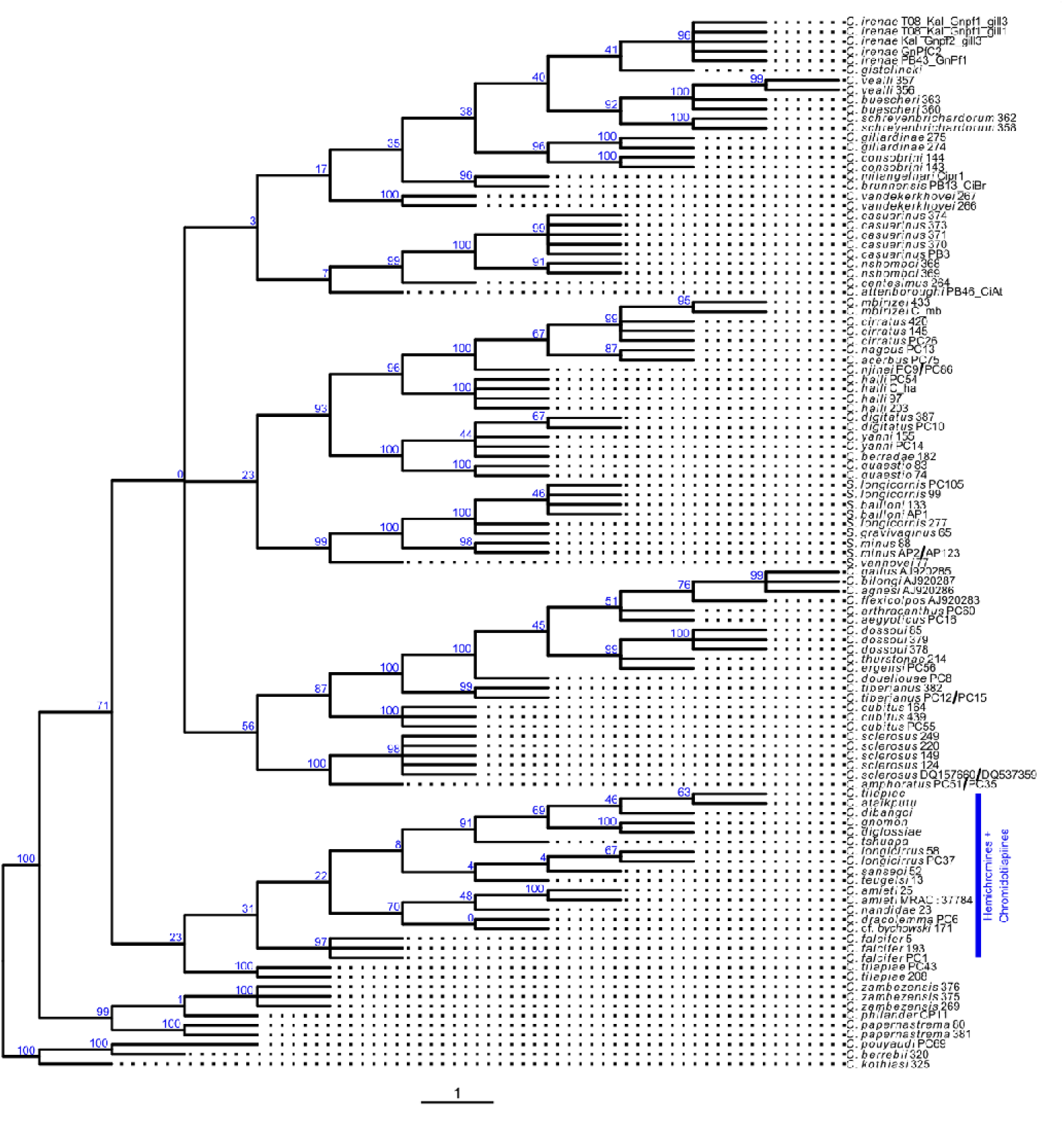
Phylogram of species of *Cichlidogyrus* inferred from morphological characters under maximum parsimony and using the molecular tree published by Cruz-Laufer et al. [12] as a constraint. Species of *Cichlidogyrus* infecting chromidotilapiines cichlids appear nested inside a species group infecting mostly hemichromine cichlids (highlighted in blue), a host tribe with a shared distribution in Central and West Africa.

The machine learning algorithms predicted clade affiliation of species of *Cichlidogyrus* with known clade affiliation with varying degrees of accuracy. Random Forest showed a moderate agreement (κ = 0.51) for species with known phylogenetic position after parameter optimization (mtry = 9) followed by artificial neural networks(κ= 0.39; size = 20, decay = 1E-04) and support vector machines (κ= 0.24; C = 3.05E-05, σ = 8192) with only fair agreements. However, new species of *Cichlidogyrus* described here were placed in different groups (*Bulb*, *EAR*, *Oreo1, Tilapiae*, *Tylo*, see [12]) but never close to species infecting hemichromine cichlids as suggested by the parsimony analysis.

## Discussion

Chromidotilapiini is one of the most species-rich tribes of cichlid fishes on the African continent, rivalled only by the hyperdiverse lineages of the Eastern African radiations and tilapias from Oreochromini [107, 111]. Despite this diversity, their relationship with other organisms in their environment remains poorly understood, especially concerning parasitic diseases. Forty years have passed since the last major parasitological studies on chromidotilapiines [74, 75] and almost 30 years since the most recent infection was reported in the literature [76]. Here, the parasite fauna of chromidotilapiines across West and Central Africa is investigated for the first time. Furthermore, this study is the first to infer the phylogenetic position of new monogenean species without molecular data using a phylogenetic analysis restricted by a baseline molecular phylogeny. Through examining the gills of specimens stored in natural history collections, ten species of dactylogyrid monogeneans were reported, of which eight are new to science, six belonging to *Cichlidogyrus* and two to *Onchobdella*.

### Monogenean evolution in Western Africa: allopatric speciation

The evolutionary history of monogenean parasites infecting cichlid fishes in West and Central Africa has been largely shaped by geographical constraints. Specifically, among the lineages of *Cichlidogyrus* and *Onchobdella* infecting chromidotilapiine, hemichromine, and pelmatochromine cichlids, we find strong indicators for allopatric speciation processes. Chromidotilapiini, Hemichromini, and Pelmatochromini are tribes of cichlids whose members occur across Central and West Africa [4,54,111]. Our study reveals that their monogenean parasites belonging to *Cichlidogyrus* and *Onchobdella* are each other’s closest relatives. Therefore, the parasite lineages have likely diverged from their relatives infecting other cichlid fishes due to the geographical isolation of the host lineages. While this pattern might also be explained by the fact that the host lineages diverged early from the haplotilapiine lineages [110, 111], chromidotilapiines, hemichromines, and pelmatochromines share no common ancestor. Therefore, co-speciation of host and parasite lineages can be excluded. The close relationship of the parasite lineages is not the result of a shared ancestry of the host species. Instead, the monogenean fauna is indicative of the shared environment where chromidotilapiines, hemichromines, and pelmatochromines occur. The incongruence of parasite and host phylogenies stands in contrast with species of *Cichlidogyrus* infecting hosts from the East African radiations, where both hosts and parasites form well-supported monophyletic clades [12, 97]. Allopatric patterns in the evolution of West and Central African monogeneans come as no surprise as this extensive region offers a multitude of ecological barriers and encompasses many isolated habitats such as river basins [53,55,120,128], lakes [62, 101], and rapids [112], all of which support high numbers of endemic fishes.

Despite their similar distribution patterns, the lineage of *Cichlidogyrus* infecting chromidotilapiines and hemichromines, and the genus *Onchobdella* (the representatives of which additionally infect pelmatochromines) are distinct groups among Dactylogyridae. Species of *Onchobdella* were first described in 1968 from species of *Hemichromis* and *Chromidotilapia guntheri* [72]. In the following decades, several species were added that were found to infect other hemichromine [3, 42] and pelmatochromine cichlids [83]. Species of *Onchobdella* have not been reported from any other host tribes despite several parasitological studies on oreochromine and coptodonine cichlids in Western and Central Africa [43,78,80,81,84,85]. Species of *Onchobdella* are also absent from all Eastern African cichlids [27,98,99] and tilapia-like cichlids across Africa [42] to our current knowledge. In contrast, species of *Cichlidogyrus* occur across Africa and the Levant [11]. Only one lineage of *Cichlidogyrus* infects the three West and Central Africa host tribes discussed here as inferred from our phylogenetic analysis (Figure 16) wherein *Cichlidogyrus ataikputu* n. sp., *C. dibangoi* n. sp., *C. ophioglossa* n. sp., *C. gnomon* n. sp., and *C. tilapiae* ex *Chromidotilapia* spp. appear as closely related to their congeners infecting hemichromine cichlids based on morphological characters of the sclerotised attachment and reproductive organs (Figure 16). These geographical distribution patterns of species of *Cichlidogyrus* and *Onchobdella* infecting hemichromines have previously been remarked upon [43]. However, the authors of these studies remained cautious about drawing any wider conclusions because of the substantial gaps concerning monogenean biodiversity in Western Africa. Among other dactylogyrid monogeneans, allopatric patterns are well-documented, such as those in European members of *Gyrodactylus* [36, 61], but the substantial lack of knowledge in terms of species numbers and distribution patterns severely limits similar research in the tropics.

Despite the results of the phylogenetic analysis, no qualitative morphological evidence of characters shared by chromidotilapiines-infecting species of *Cichlidogyrus* and their hemichromine-infecting congeners (*Hemi* clade, see [12]) was found. For example, the looped or spiralled copulatory tube typical for the members of the *Hemi* clade, represents a feature that is absent from the chromotilapiine-infecting species. Such discrepancies may also explain the ambiguous nature of the results obtained through the machine learning analysis across all algorithms employed in this study. Furthermore, few morphological similarities were revealed between chromidotilapiines-infecting species. *Cichlidogyrus ophioglossa* n. sp. and *C. gnomon* n. sp. share a distal plate that is associated with the accessory piece of the MCO. *Cichlidogyrus ataikputu* n. sp.*, C. dibangoi* n. sp., and *C.* ‘*Pelvicachromis roloffi*’ have a simple *bauplan* (a mostly straight accessory piece and copulatory tube), reminiscent of the “tilapia” parasite *C. tilapiae*. Finally, the copulatory organ morphology of *C. tshuapa* n. sp. is reminiscent of the spiralling accessory piece of *C. papernastrema*, perhaps suggestive of a link to a group of southern African species including *C. philander* Douëllou, 1993 and *C. zambezensis* Douëllou, 1993 [12]. However, previous studies indicate that some morphological similarities in the attachment and reproductive organs of dactylogyrid monogenean may in fact be a result of convergent evolution, such as the marginal hook length in species of *Cichlidogyrus* [12], the retention of ancestral features, e.g. the characters of the polyphyletic genera *Demidospermus* Suriano, 1983, *Haliotrema* Johnston & Tiegs, 1922, and *Ancyrocephalus* Creplin, 1839 [18,48,49], or host switching, e.g. hook lengths in *Cichlidogyrus amieti* Birgi & Euzet, 1983 [66]. Checking for such potential contradictions between morphological patterns and subsequent molecular-phylogenetic results, once genetic data are available, is an important research target for future studies. In particular, congruence analyses [49] or phylogenetic comparative methods [12,46,104] as applied by previous studies might provide further insight into the evolutionary history of this lineage of *Cichlidogyrus*.

### One host, several parasites: intra-host speciation and host switching

Alongside the discussed allopatric mechanisms, several instances of host sharing of species of *Cichlidogyrus* and *Onchobdella* are revealed in the current study. Host sharing can result from intra-host speciation and host switching. Recent publications indicate that parasites undergo cycles of niche isolation (e.g. intra-host speciation) and expansion of host repertoires (e.g. host switches) [8]; this fluctuation is also considered a likely occurrence in monogenean flatworms [6, 10]. However, identifying patterns for intra-host speciation or host switching in the absence of DNA sequence data poses a major challenge. In the present case, the differentiation in the attachment organs might signal an adaptation to specific microhabitats similar to reports from Europe on species of *Dactylogyrus* and *Lamellodiscus* Johnston & Tiegs, 1922 [88, 116]. For instance, *Onchobdella macrohamulus* n. sp. has a much larger set of dorsal anchors than the co-infecting *O. yemojae* n. sp. or any other species of *Onchobdella*. This difference might suggest an adaptation to a different gill microhabitat, where larger sclerites with more leverage (see [103]) are required, but this feature may equally be an ancestral character from a separate lineage of *Onchobdella*. No species with a similar morphology has been found to date and currently DNA sequences for only two species of *Onchobdella* are available [65]. This lack of data means that the phylogenetic relationships of species of *Onchobdella* remain currently in obscurity. Similar questions to those for *Onchobdella* also arise for co-occurrences of species of *Cichlidogyrus* on chromidotilapiines, although, in this study, the species found on chromidotilapiines appear to form a monophyletic group (Figure 16). Co-infections of members of *Cichlidogyrus* with members of *Onchobdella* are similar to co-infections of these same groups reported for hemichromine and pelmatochromine cichlids [43, 83]. *Onchobdella* and *Cichlidogyrus* form part of Dactylogyrinae *sensu* Kmentová & Cruz-Laufer et al., [49], a subfamily of Dactylogyridae, but are otherwise unrelated. Niche specialisation may represent a strategy to avoid competition and to facilitate co-infections of closely-related monogeneans [105,106,115] resulting in microhabitat preferences [26]. For instance, host sharing between species of *Cichlidogyrus*and *Onchobdella* may be indicative of distinct ecological niches on the hosts’ gills, thereby enabling the co-existence of the two lineages. A detailed analysis of the gill microhabitats such as in [26, 116] should be implemented so as to provide more insight into the niche habitats of these monogenean species.

### No speciation or cryptic speciation?

The crater lakes in Cameroon have long been of interest to evolutionary biologists as they represent not only the location of one of the most prominent examples of sympatric speciation, but are also home to many endemic species [5,62,69,101]. Species of *Coptodon* and *Sarotherodon* have formed four species flocks in a total of some 25 species in lakes Bermin, Ejagham, and Barombi Mbo [62]. In the present study, the gills of specimens of *C. guntheri loennbergi*(Trewavas, 1962), which is endemic to Lake Barombi Kotto were also screened. Unlike the other lakes, Barombi Kotto supports only few species, of which *Coptodon kottae* (Lönnberg, 1904) and *C. guntheri loennbergi* are the only endemic examples [5, 122] although, the status of the latter—*C. guntheri loennbergi—*has been contested due to the absence of apparent morphological delimiters [54]. Monogeneans have shorter generation time than their hosts, making them a more efficient subject of study when differentiating host population structure (*magnifying glass effect*, see [21]). Yet *Cichlidogyrus ataikputu* n. sp. and *C. dibangoi* n. sp. occur both in the lake and the surrounding river systems suggesting a lack of differentiation. Therefore, our observations provide additional evidence that *Chromidotilapia guntheri loennbergi* should be considered a synonym of *C. guntheri*.

Despite the morphological similarities used to described each species, the presence of cryptic species is also possible among monogeneans and species of *Cichlidogyrus*. For example, *Cichlidogyrus halli* has previously been identified as a species complex as it displays substantial morphological variation [16, 42]. It was only through a detailed analysis that morphological and genetic differences could be highlighted between specimens of *Cichlidogyrus halli* sensu stricto from Zimbabwe leading to the discovery of a new species [22]. High variation in the DNA barcodes suggest that similar issues apply to several species originating from the Congo Basin [40]. One potential species complex in the present study is *Cichlidogyrus ophioglossa* n. sp., for which a string-like structure in the MCO shows considerable structural variation (Figure 4). A similar case has been reported for *C. cirratus* Paperna, 1964, where a thin transparent structure in the MCO varies considerably. This variation was initially associated with the position of the worm on the slide [132], but recent evidence suggests the presence of three distinct morphotypes [60]. For *C. ophioglossa* n. sp., molecular phylogenetic analyses using markers with a high evolutionary rate (e.g. the internal transcribed spacers or the cytochrome oxidase *c* subunit I gene) may provide valuable insights into cryptic diversity. Furthermore, scanning electron microscopy or confocal microscopy can be used to obtain three-dimensional images of complex structures such as the MCO, which will provide a better insight in their morphology as well as decreasing the probability of sample destruction [19,109,132].

A further example for a potential species complex is *Cichlidogyrus tilapiae* [93] characterized by a relatively simple *bauplan* of its MCO with a straight copulatory tube and accessory piece with no remarkable protruberances other than a slight distal hook. Some species described in the current study strongly resemble *C. tilapiae,* including *C. ataikputu* n. sp. and *C. dibangoi* n. sp. In point of fact, several specimens were found belonging to *Cichlidogyrus* (*C. tilapiae* ex *Chromidotilapia* spp.) (Figure 15) that are morphologically indistinguishable from *C. tilapiae* despite our parsimony analysis placing these specimens in a separate lineage together with the other species of *Cichlidogyrus* described herein (Figure 16). This result is also confirmed by the PCA (Figure 14), where measurements of *C. tilapiae* reported from other hosts in previous publications [12, 100] form a cluster distinct from the specimens reported here. However, no qualitative characters were found that delineate the specimens infecting chromidotilapiine cichlids as a separate species. The relationships in this species complex may be resolved only through detailed morphological and molecular studies of the target taxon.

### Conclusive remarks: natural history collections and morphological data

The present manuscript represents the most extensive study on monogeneans infecting Western and Central African fishes based solely on historic host collections to date. However, the two methods employed to identify the phylogenetic position of the newly described species of *Cichlidogyrus* (parsimony and machine learning analyses) showed varying levels of success. While the parsimony analysis provided results indicative of the well-supported hypothesis that chromidotilapiine-infecting species are closely related to their congeners infecting hemichromine cichlids, the machine learning approach that showed promise in a recent publication [12] failed to provide any conclusive results despite the use of multiple algorithms and additional morphological characters. For the time being, we recommend that the use of these algorithms be reserved as a means to quantify the predictive power of characters rather than serving to approximate phylogenetic positions.

As the biodiversity of metazoan parasites remains vastly underexplored, the present study clearly demonstrates that collection-based studies of ectoparasites are an effective tool for describing the parasite fauna of rare hosts, despite the absence of high-quality DNA samples. Recent studies also highlight the fact that these collections provide windows into the past in terms of human-induced changes of host-parasite communities [41, 129]. Morphological and collection-based studies of these organisms, therefore, unequivocally remain an essential part of taxonomic exploration.

## Acknowledgements

We thank Miguël Parrent and Jos Snoeks for providing access to the ichthyological collection of the Royal Museum for Central Africa (RMCA). We thank Nnoye Scholastica Mbanu and Jennifer Mbanu for suggesting species epithets in Igbo. Part of the research leading to results presented in this publication was carried out with infrastructure funded by the European Marine Biological Research Centre (EMBRC) Belgium, Research Foundation – Flanders (FWO-Vlaanderen) project GOH3817N. AJCL (BOF19OWB02), NK (BOF21PD01) and MPMV (BOF20TT06) received support from the Special Research Fund of Hasselt University.

